# Extracellular histones, a new class of inhibitory molecules of CNS axonal regeneration

**DOI:** 10.1101/365825

**Authors:** Mustafa M. Siddiq, Sari S. Hannila, Yana Zorina, Elena Nikulina, Vera Rabinovich, Jianwei Hou, Rumana Huq, Erica L. Richman, Rosa E. Tolentino, Jens Hansen, Adam Velenosi, Brian K. Kwon, Stella E. Tsirka, Ian Maze, Robert Sebra, Ravi Iyengar, Marie T. Filbin

**Author notes:** Co-Senior Authors (Ravi Iyengar, Marie T. Filbin). Deceased January 15, 2014 (Marie T. Filbin). Address correspondences to: Ravi Iyengar Dept of Pharmacological Sciences, Box 1215, 1425 Madison Ave Rm 12-70 New York NY 10029 and Mustafa M. Siddiq.

## Abstract

Axonal regeneration in the mature CNS is limited by extracellular inhibitory factors. Triple knockout mice lacking the major myelin-associated inhibitors do not display spontaneous regeneration after injury, indicating the presence of other inhibitors. Searching for such inhibitors we have detected elevated levels of histone H3 in human cerebrospinal fluid (CSF) 24 hours after spinal cord injury. Following dorsal column lesions in mice and optic nerve crushes in rats, elevated levels of extracellular histone H3 were detected at the injury site. Similar to myelin-associated inhibitors, these extracellular histones induced growth cone collapse and inhibited neurite outgrowth. Histones mediate inhibition through the transcription factor YB-1 and Toll-like receptor 2, and these effects are independent of the Nogo receptor. Histone-mediated inhibition can be reversed by the addition of activated protein C (APC) *in vitro*, and APC treatment promotes axonal regeneration in the crushed optic nerve *in vivo*. These findings identify extracellular histones as a new class of nerve regeneration-inhibiting molecules within the injured CNS.

**One sentence summary:** Proteins typically associated with chromatin structure play an unexpected role in limiting axonal regeneration after injury.

## Introduction

Axonal regeneration following injury is not well understood. Hence development of therapeutic strategies for neurorepair are challenging. Central nervous system (CNS) myelin proteins and chondroitin sulfate proteoglycans (CSPGs) are well-characterized inhibitors of axonal growth (1-4), but genetic and pharmacological elimination of these factors elicits only a modest increase in axonal regeneration following spinal cord injury (5). This suggests that other inhibitory molecules are present in the CNS environment. Extracellular histones have been detected in neurodegenerative diseases such as Alzheimer’s disease, scrapie, and Parkinson’s disease (6-8), and histone H1 has been shown to induce microglial activation and neuronal apoptosis. It has also been reported that histones are released from apoptotic and necrotic cells following cerebral ischemia, lung injury, and myocardial infarction (7, 9). Previous studies on sepsis have shown deleterious effects of extracellular histones as they stimulate apoptotic pathways to trigger cell death (6). As ischemia and cell death are key pathophysiological events in axonal injury, we hypothesized histones could be released from damaged cells and contribute to the pathophysiology by inhibiting axonal regeneration. To test our hypothesis, we determined if histone levels are elevated in humans after spinal cord injury. We show similar effects after spinal cord in injury and optic nerve crush in murine models.

Using cell-based assays for axonal regeneration we show that histones inhibit outgrowth through TLR2 and the transcription factor YB1. We also show that the protease APC that selectively cleaves histones (6) and enhance neuroprotection in cerebral ischemia (10) relieves histone induced inhibition of axonal outgrowth. In vivo APC promotes regeneration of crushed optic nerves. Histones have been shown to bind receptors such as Toll-like receptor 2 and 4 (TLR2 and TLR4), outside the nervous system (11). Histones binding to TLR2 and 4 can activate intracellular pathways in those tissues (12). In this study, we demonstrate that histones are elevated in response to spinal cord injury (SCI) in humans. Using a rat model we show that histones inhibit neurite outgrowth in vitro, and that these effects are mediated through TLR2 and the transcription factor YB-1. Furthermore, we show that APC promotes regeneration of the rat optic nerve in vivo.

## Results

### Detection of histones in the CSF of humans with SCI

We obtained CSF from humans with SCI with an ASIA Grade A (complete paralysis), approximately 24 hours post-injury from the International Spinal Cord Injury Biobank (ISCIB). From the 3 human specimens, all had Thoracic injury at T6, T10 and T4. We also have 4 controls from humans that are normal (ASIA Grade E). We ran the CSF samples on a gradient gel and probed for human H3 histones (Fig. 1A), the individual running the gel was blinded to what the samples were. We were able to detect the histones in the CSF of the human specimens with spinal cord injury. None of the control normal human CSF had detectable levels of H3. We normalized all the samples to albumin in the CSF (Fig. 1B).

**Figure 1.**
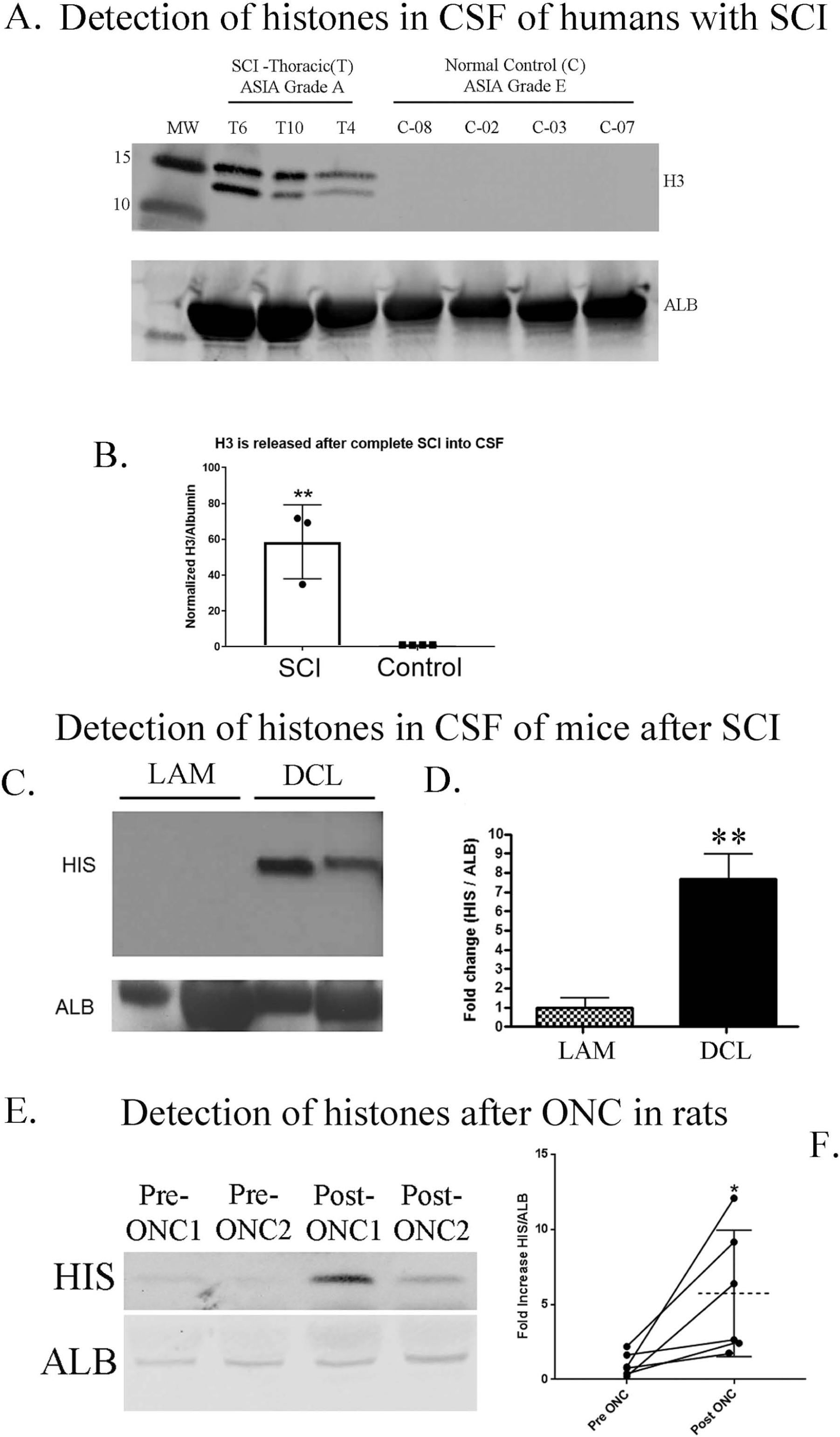
Elevated levels of extracellular histones are detected in the injured CNS. **A**. From human CSF collected from patients approximately 24 hrs post-injury with ASIA score of A (complete impairment, paralysis), and the controls are all normal (ASIA Grade E). On a gradient western blot, we loaded 30µgs of sample for each human specimen and probed for histone H3 or Albumin for normalizing control. The H3 is at the predicted size of 17kDa and we detect it in patients with Thoracic injury (T6,T10 and T4). **B.** The bar graph is SCI patients comparing to controls, where we see a very significant **p<0.01 by t-test. Elevated levels of Histone H3 (HIS) in the CSF fluid collected from mice with DCL are detected by western blot, compared to laminectomy (LAM) alone. **C.** Histone H3 levels are normalized to Albumin (ALB) and in the bar graph is the average of 5 animals (n=5 for both DCL and LAM group) with DCL where we see a significant elevation of free Histone H3 in the CSF fluid. **D**. Adult rats first had their optic nerves exposed and we applied gelfoam over the uninjured nerve for 48hrs (Pre-ONC1&2), subsequently removing the gelfoam into PBS with protease inhibitor cocktail, we crushed the optic nerve and applied a fresh piece of gelfoam over the injury site. We removed the gelfoam 48hrs later (Post-ONC 1&2). We found within the same animal a significant increase in HIS levels after crushing the optic nerve (Post-ONC) than prior to injury (Pre-ONC), data was graphed in **F**. (n=6, see Table 1 for full comparison of Pre- and Post-ONC levels of HIS). All statistics were performed by t-test, *p<0.05 and **p<0.01, except Figure F. paired t-test.

**Table 1.** Elevated levels of histones-released into gelfoam after ONC in the same animal. Below we show the band intensity detected by western blotting for Histone H3 from the gelfoam placed before and after optic nerve crush, in the same animal.

| **Animal#** | **Pre-ONC** | **Post-ONC** | **Fold increase** |
| --- | --- | --- | --- |
| 1 | 0.21183 | 6.385752 | 30.15 |
| 2 | 0.827125 | 12.08353 | 14.61 |
| 3 | 0.75167 | 1.755859 | 2.34 |
| 4 | 0.384941 | 2.419942 | 6.29 |
| 5 | 2.198021 | 9.15946 | 4.17 |
| 6 | 1.626405 | 2.637283 | 1.62 |

To validate this finding in the human specimens with Thoracic (T) injury, we performed T8-9 level dorsal column lesions (DCL) in 9-12 weeks old 129S1/SvImJ mice (The Jackson Laboratory) and collected cerebrospinal fluid (CSF) from the injury site 24 hours later. When compared to a sham group that received laminectomy (LAM) alone, histone levels were significantly higher in CSF obtained from spinal cord-injured animals (Figs. 1C&D). To provide further evidence of histone release after CNS injury, we exposed the optic nerve in adult rats and placed gelfoam on the uninjured nerve. These pre-injury gelfoam samples were extracted 48 hours later. The nerve was then re-exposed and crushed and a new piece of gelfoam was placed at the injury site for 48 hours, enabling us to compare histone levels pre-and post-injury in the same animal. Western blot analysis revealed that histone levels were visibly increased in the optic nerves of individual animals following injury, and overall, this increase was statistically significant (Figs. 1E&F). Quantification of the histones released into the gelfoam confirmed that histone levels within individual animals (n=6) were significantly higher following optic nerve crush (ONC; Table 1).

### mRNA-seq analysis

We also performed bulk mRNA-Seq experiments comparing control (non-treated) primary rat cortical neurons to neurons treated for 2hrs with a mixed population of histones isolated from calf thymus [10μg/ml]. The full list of Differential Expressed Genes (DEGs) are found in Table 2. Analysis of the subcellular pathways (SCPs) using WikiPathways revealed that one of the top 10 up-regulated SCPs was for Spinal Cord Injury and 4 out of the 7 down-regulated pathways found by Geneontology (GO) involve axon development (Figs. 2A-C). This supports the role of histones in mediating inhibition of neurite outgrowth.

**Figure 2.**
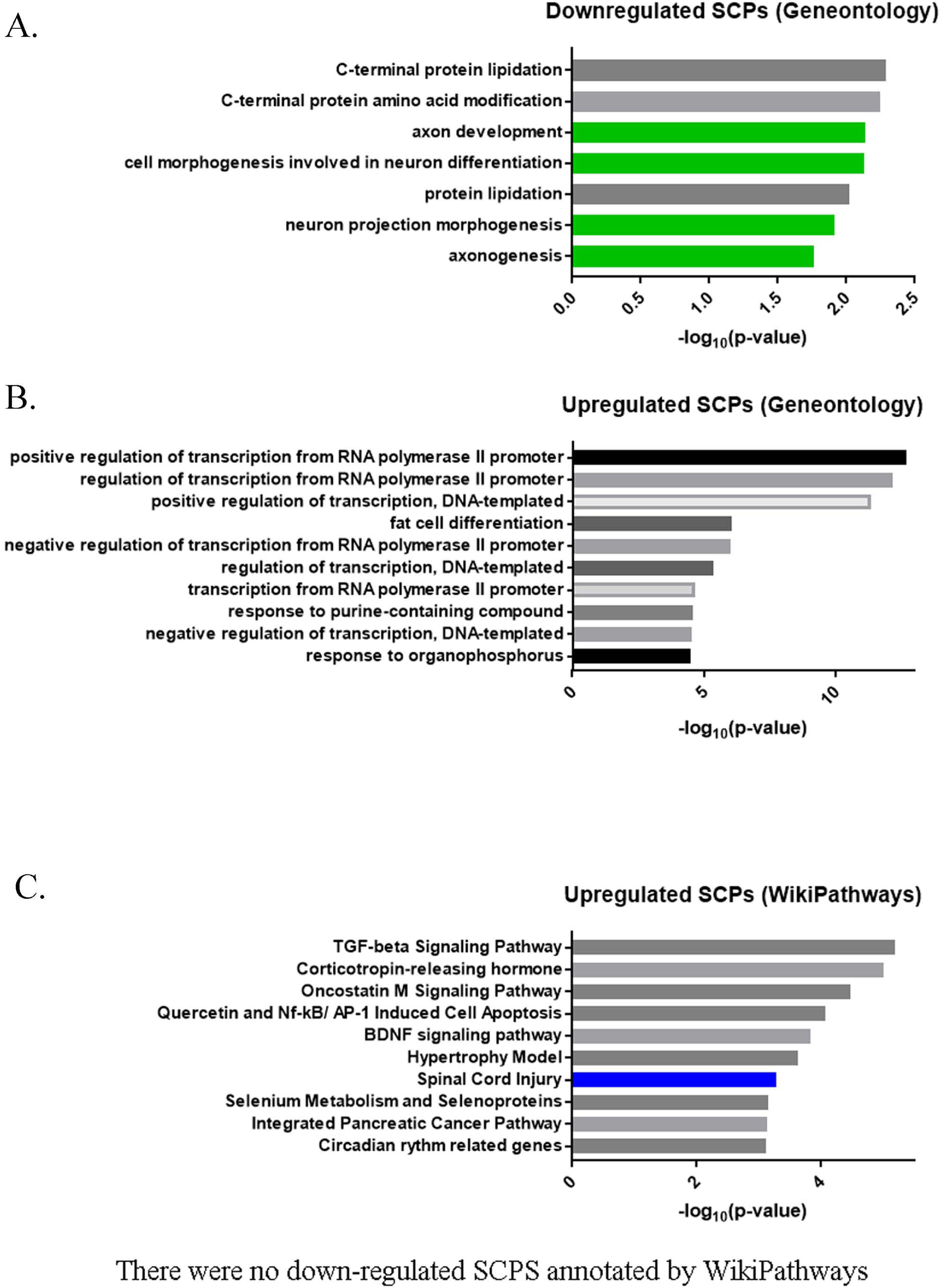
bulk mRNA-seq analysis with cortical neurons treated with histones. **A.** Up- and Down-regulated genes for histone treated cortical neurons were subjected to pathway enrichment analysis to identify top-ranked pathways. From Geneotology (GO), the top-ranked (according to P values) down-regulated SCPs related to histone treatment of cortical neurons and **B.** up-regulated SCPs annotated by GO. **C.** From Wikipathways, the top-ranked (according to P values) up-regulated SCPs related to histone treatment of cortical neurons, there were no down-regulated SCPs related to histone treatment annotated by Wikipathways.

**Table 2.**
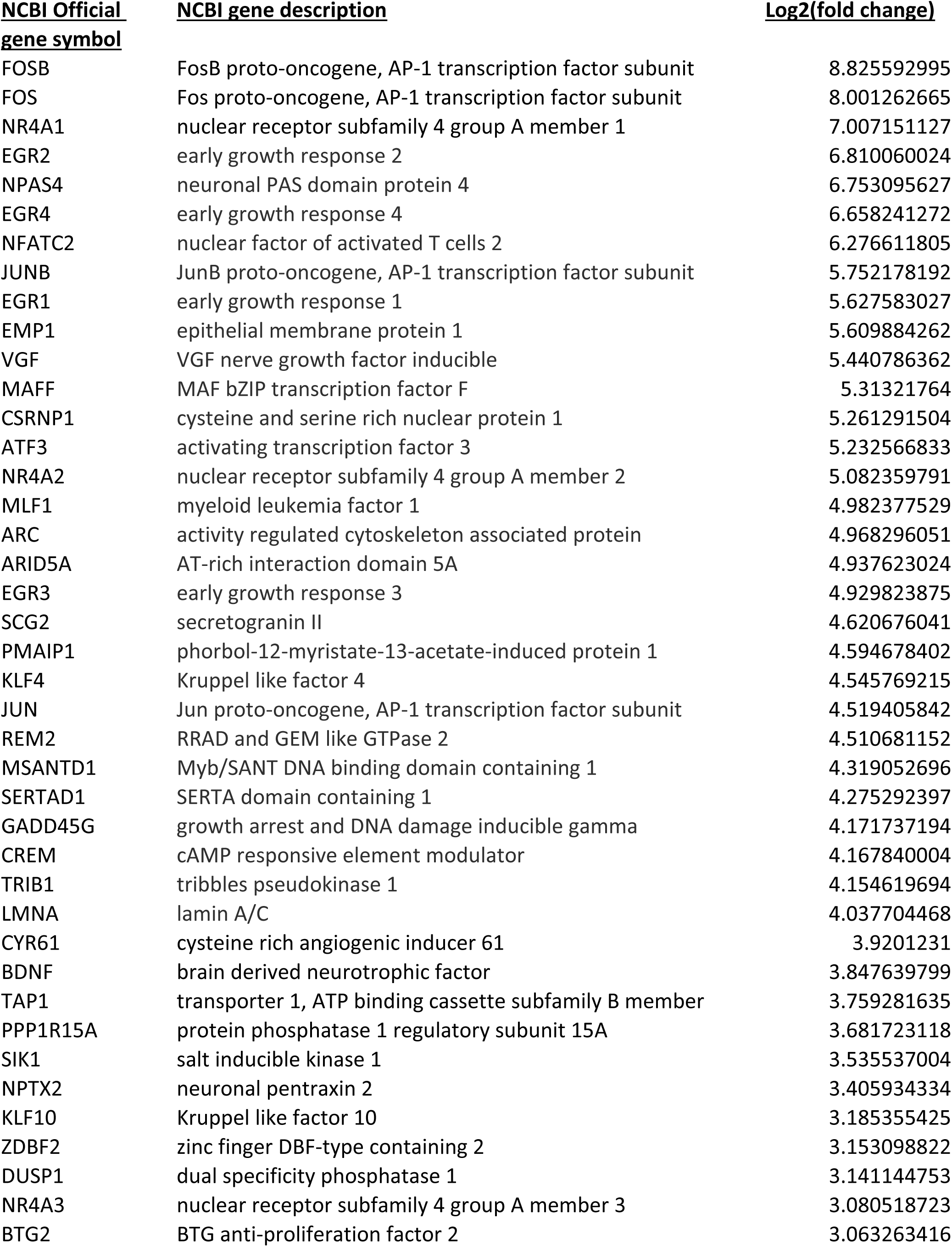
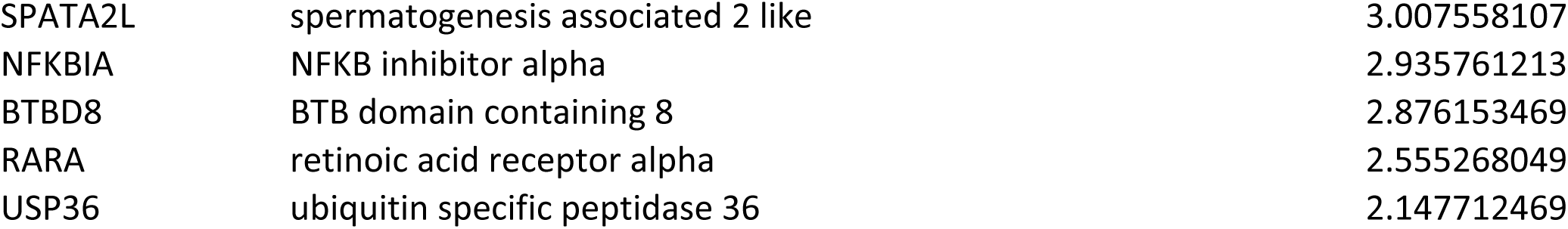

Studies of myelin-associated inhibitors and CSPGs have shown that these molecules inhibit axonal regeneration through activation of the small GTPase RhoA, which mediates growth cone collapse (13). To examine the effects of histones on growth cone morphology, we treated cortical neurons with aprotinin (APN) or histones and then immunostained growth cones for β-III tubulin and actin. Cortical neurons treated with aprotinin, a protein of similar size and isoelectric point to histones (but with no structural homology), did not affect growth cones as their morphology was similar to those seen in non-treated cortical neurons (Fig. 3A). But within 10 minutes of treatment with histones (10μg/ml), we observed more actin at the growth cone tips, and this becomes very apparent within 15 minutes (Fig. 3A). Furthermore, the β-III tubulin shows a punctate pattern of staining, suggesting degeneration. By 25 minutes, dysmorphic end-bulbs are apparent and these are still present at 14 hours. With regard to the molecular events underlying these morphological changes, we observed that a 30 minute treatment with extracellular histones significantly increased activation of RhoA (Fig. 3B) and also reduced levels of p35 (Fig. 3C), a protein that can overcome MAG-mediated inhibition by activating cyclin-dependent kinase 5 (14). This provides evidence that the signaling pathways activated by CNS myelin, CSPGs, and histones converge at key points to mediate inhibition, and importantly, it also shows that histones can affect cytoskeletal dynamics. To assess the effects of extracellular histones on axonal growth, we then performed a series of neurite outgrowth assays. Using permissive substrates of Chinese hamster ovary (CHO) cells, we observed that histones inhibited neurite outgrowth in a dose-dependent manner for both cortical (Figs. 3D for graph and S1A-C for representative images of cortical neurons on CHO cells) and dorsal root ganglion neurons (Fig. S2). We performed a second neurite outgrowth assay using a permissive layer of primary rat astrocytes. Neurite outgrowth was significantly reduced in a dose-dependent manner in response to histone treatment (Figs. 3E&F). This effect was not due to toxicity or the presence of other inhibitory factors, as the astrocyte monolayers were unaffected by the histones (Figs. 3G), and histones did not induce expression of the CSPGs neurocan and brevican in astrocytes (Fig. S3). Finally, to elucidate the contributions of specific core histones to inhibition, cortical neurons were plated in microfluidic chambers, and recombinant histones H3 and H4 were applied to the neurite compartment. Both histone H3 and H4 inhibited neurite outgrowth, with histone H3 having a more potent effect (Fig. 3H, full chambers are in Fig. S5). Morphologically, neurites treated with either histone H3 or H4 displayed the prominent end bulbs typical of dystrophic axons (Fig. 3I and S4). Quantification of the neurite outgrowth in Fig. 3H is in Fig. 3J.

**Figure 3.**
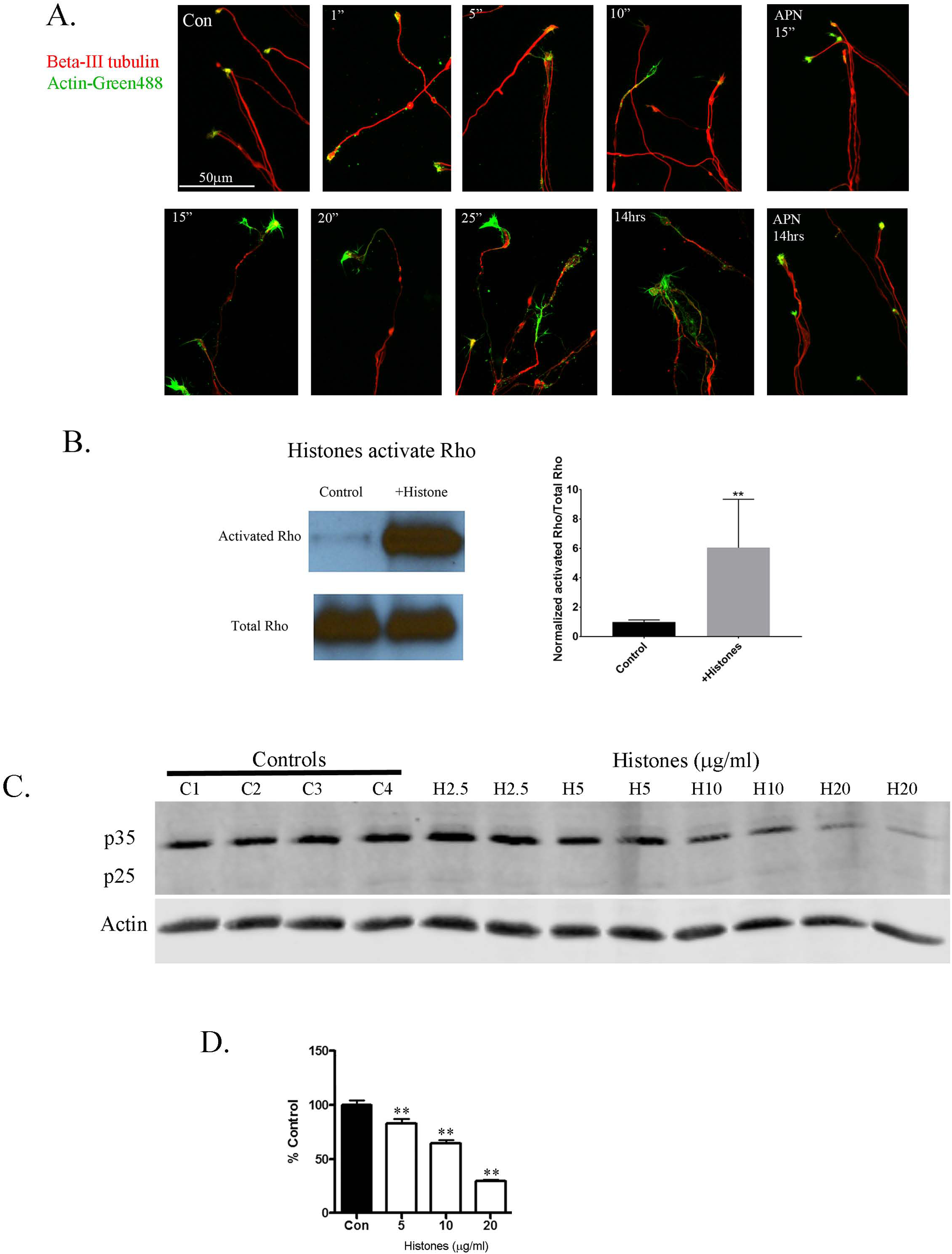
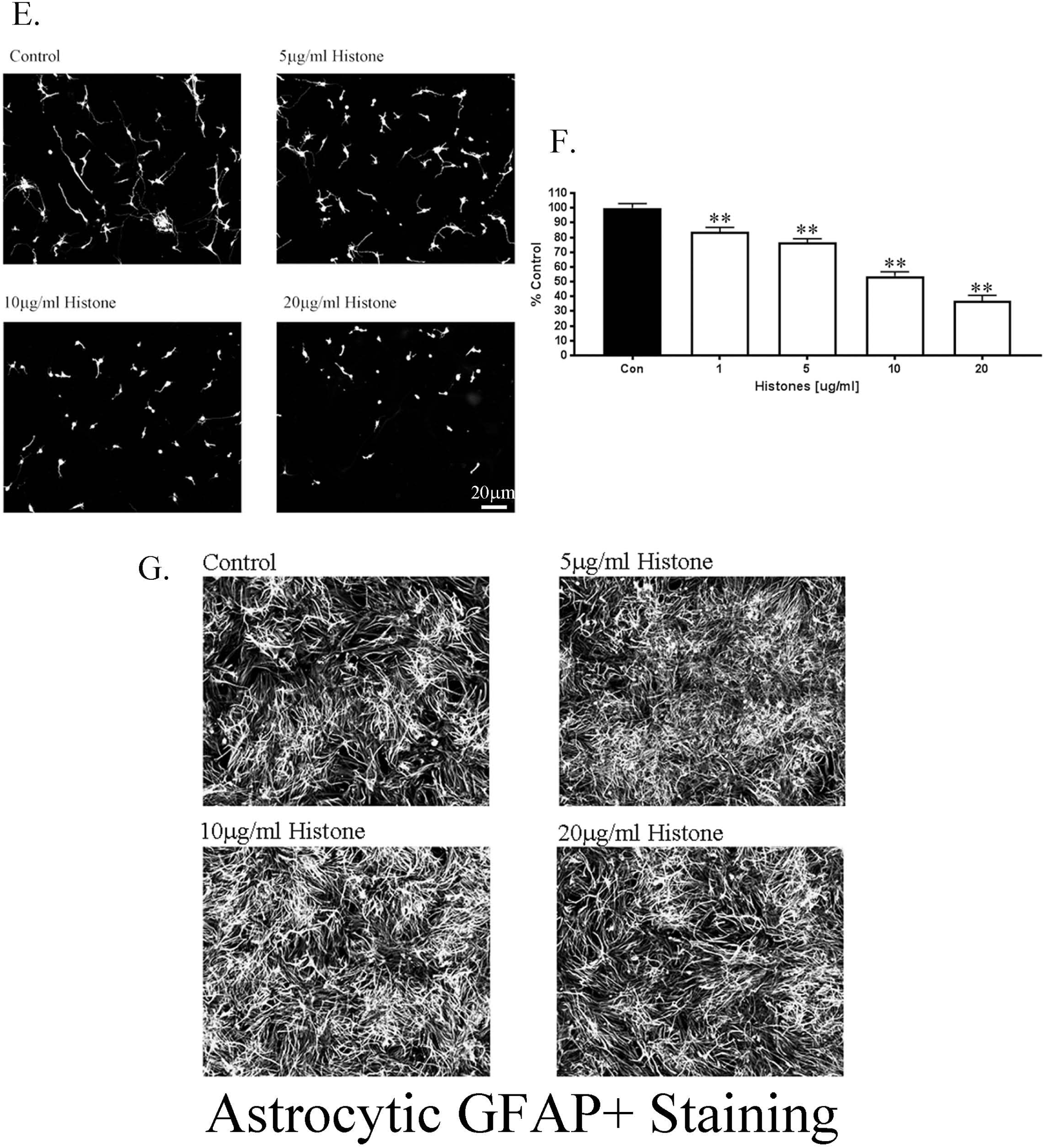
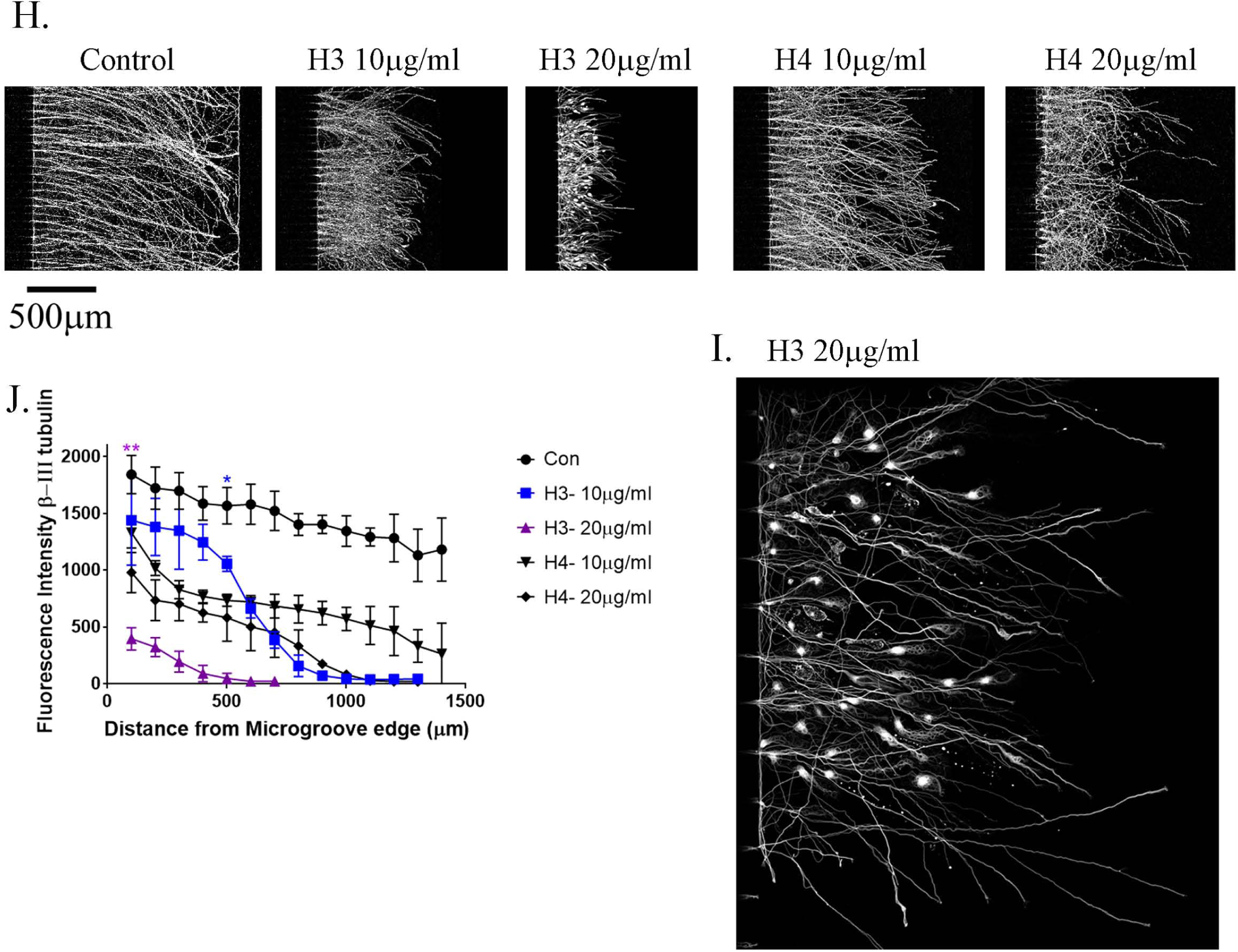
Extracellular Histones are inhibitory to neurite outgrowth. **A.** Cortical neurons on PLL-coated glass plates. Images of β-III tubulin in red and actin in green, displaying growth cones, imaged at 63X. Control, non-treated or Aprotinin (APN) for 15mins or 14hrs has no effect on growth cone morphology. Histones applied (1, 5, 10, 15, 20, 25 mins or 14hrs) results in dystrophic looking bulbs within 15mins. **B.** Histone applied to cortical neurons induces activation of Rho GTPase, the graph is the average (+/- SD) of 5 experiments. **C.** Histone applied to cortical neurons results in dose dependent decrease (2.5-20µg/ml) in p35 levels without any detectable p25. **D.** Cortical neurons on a permissive CHO monolayer put out long neurites (black bars, Con) but in the presence of histones they put out significantly shorter neurites (white bars). **E.** Primary rat cortical neurons on an astrocytic monolayer put out long neurites (Control), but in the presence of increasing concentrations of a mixed population histone preparation they put out dose-dependent shorter neurites as determined by β-III tubulin staining. **F.** Primary rat cortical neurons on an astrocytic monolayer put out long neurites (black) as quantified by longest length by Metamorph analysis, but there is a dose-dependent decrease in neurite length in the presense of histones. This bar graph is the average of three independent experiments with a minimum of N=300 neurons per group. **G.** Representative images of GFAP+ astrocytic monolayers in the absence or presence of 5-20µg/ml as indicated. **H.** Using PLL-coated microfluidic chambers, cortical neurons grow long neurites across the 450µM microgroove as detected by β-III tubulin. Treating the neurite growing compartment with either recombinant H3 or H4 Histones resulted in significantly shorter neurites, with H3 having a more potent effect, as seen in close-up image of H3 with 20µg/ml H3 in **I**. **J**. We quantified the neurite outgrowth in the chambers using Image J, each treatment is the average of three independent experiments. The purple ** at 100microns is for H3 20µg/ml, all points from 100-700 microns were p<0.01 compared to controls. The blue * at 500microns is for H3 10µg/ml, when it is significantly shorter compared to controls. All statistics were performed by ANOVA or t-test, *p<0.05 and **p<0.01.

In addition, we used a mixed population of histones isolated from calf thymus and showed a dose-dependent decrease in neurite outgrowth (Fig. 4A, full chambers are in Fig. S6). Aprotinin-treated neurons were similar to non-treated cortical neurons, which showed no difference in neurite outgrowth (Fig. 4A). Since aprotinin (APN) has no effect on neurite outgrowth we conclude that the inhibition that is observed was histone-mediated, and not due to nonspecific steric hindrance. Previous studies in other organ systems have reported that the deleterious effects of extracellular histones can be attenuated by activated protein C (APC), a coagulation factor protease that has been shown to specifically cleave histones H3 and H4 (6), and enhance neuroprotection in cerebral ischemia (10). Using cortical neurons plated in microfluidic chambers, we observed that administration of APC blocked histone-mediated inhibition of neurite outgrowth (Fig. 4B, full chambers are in Fig. S7).

**Figure 4.**
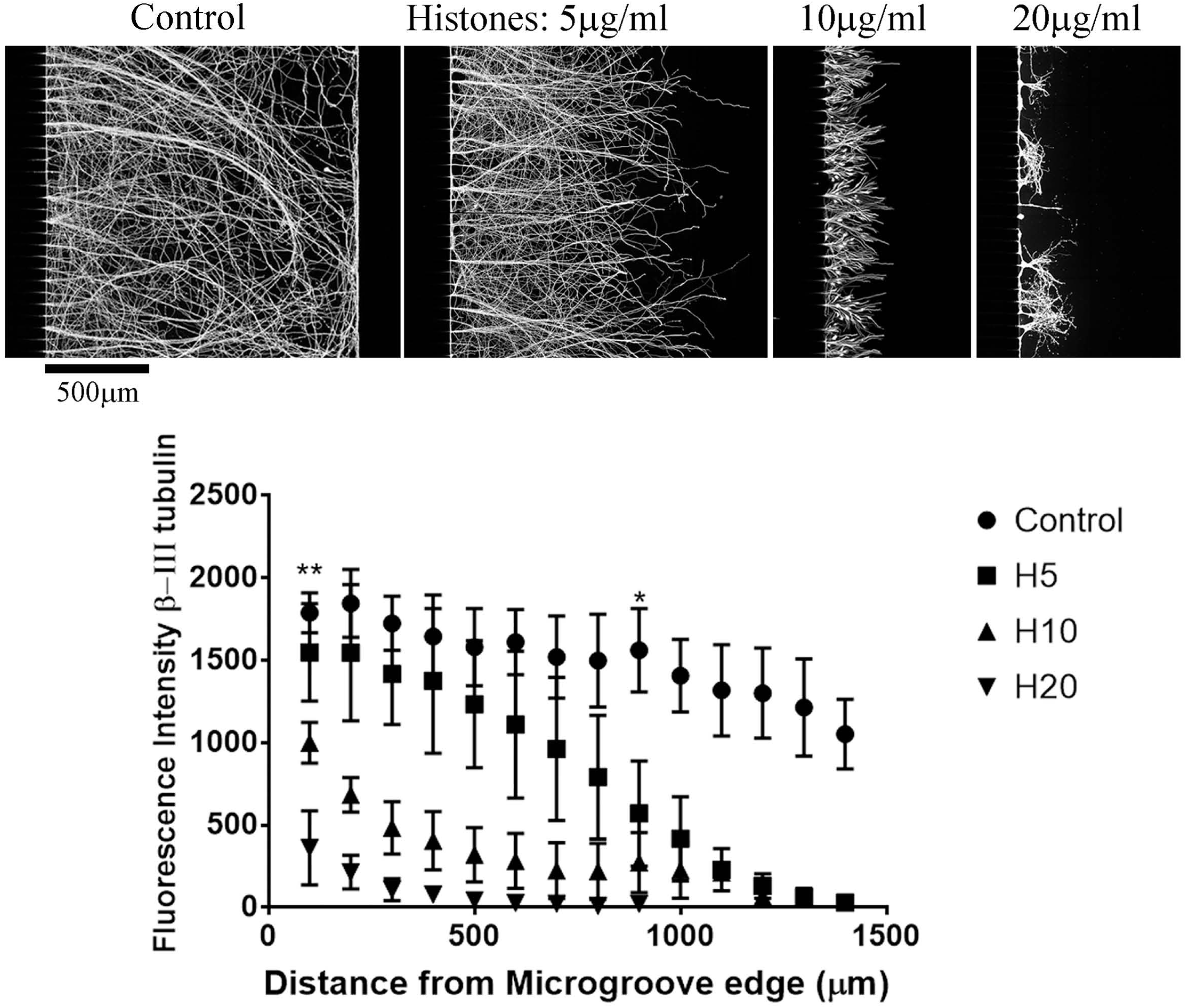
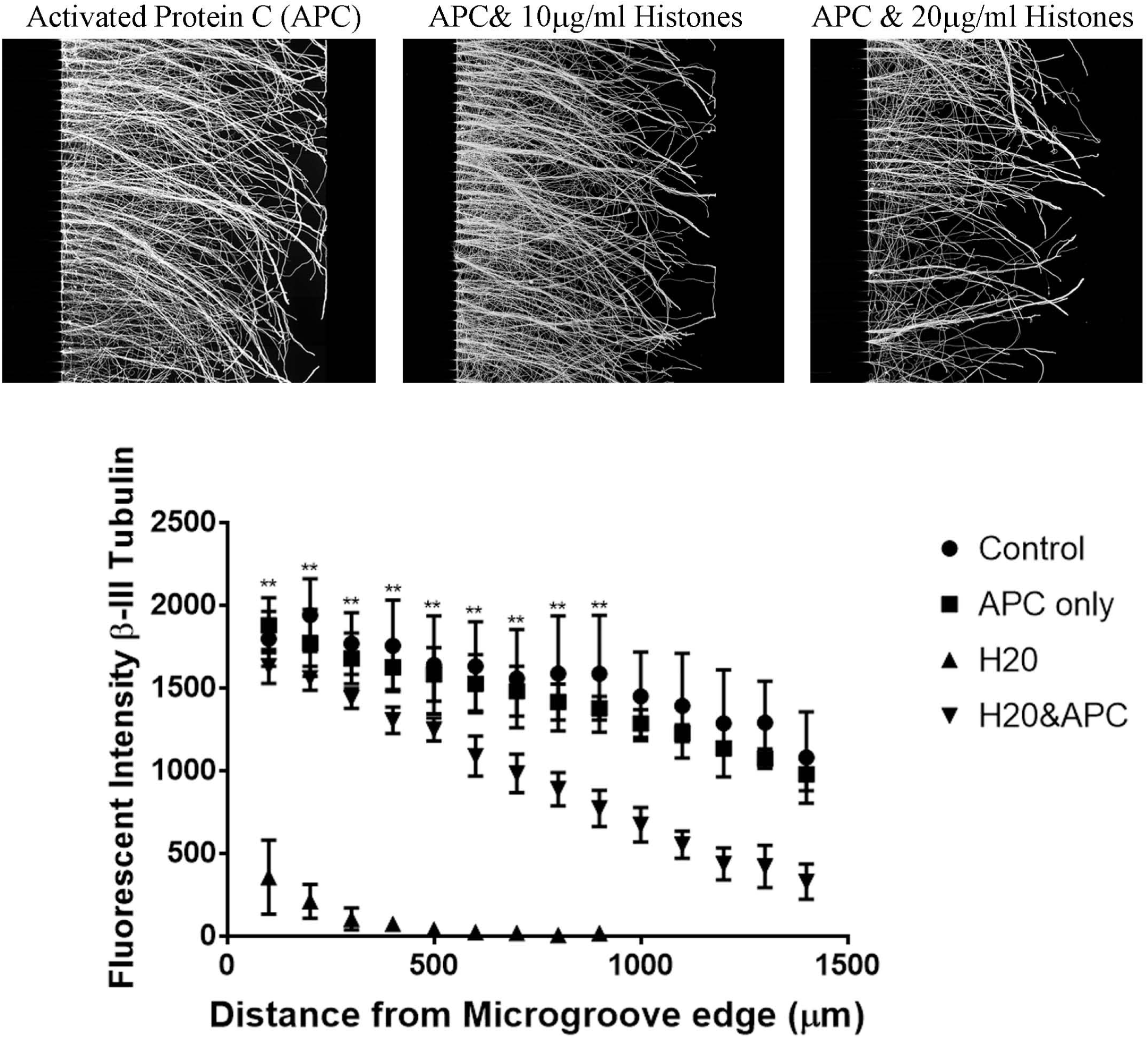
Activated Protein C (APC) blocks the inhibitory effect of Histones and promotes axonal regeneration after ONC. **A.** Cortical neurons plated on PLL coated microfluidic chambers grow robustly across the 450µm microgroove, determined by β-III tubulin staining. Treating the neurite growing side (right side from the microgroove) only with increasing concentrations of mixed population of histones isolated from calf thymus results in significantly shorter neurites. In the graph we show the average of four independent experiments, we have measurements of β-III tubulin fluorescence in increments of every 100microns from the microgroove with Control (Aprotinin-treated), H5 (Histones - 5µg/ml), H10 (Histones 10(µg/ml)) and H20 (Histones 20(µg/ml)). For statistics, the ** at 100 microns is for H10 and H20 being significantly shorter than controls. At 900 microns, H5 is significantly shorter (*) compared to controls. **B.** APC alone has no effect on neurite outgrowth, combining APC with histones and applying to the neurite side reverses the inhibitory effect of histones, restoring long neurite outgrowth. In the graph we show the average of three independent experiments, we have measurements of β-III tubulin fluorescence in increments of every 100microns from the microgroove with Control (non-treated), APC only (APC-treated), H20 (Histones 20(µg/ml)) and H20 (Histones 20(µg/ml) with APC). For statistics, the ** is comparing H20 to H20 with APC.

To elucidate the molecular mechanism underlying histone-mediated inhibition of neurite outgrowth, we treated cortical neurons with histones for 24 hours and then plated them onto permissive CHO cell monolayers in the absence of histones (18). Neurite outgrowth was severely impaired in these neurons (Fig. S8), which suggested that their intrinsic capacity for neuritogenesis had been altered at the molecular level by the histones. To determine if this effect involved the activation of transcription factors, we incubated cortical neurons with extracellular histones for 30 or 120 minutes and analyzed the lysates using a Panomics Transcription Factor protein array (19). This assay revealed significant increases in eukaryotic Y-box-binding protein1 (YB-1) in histone-treated neurons (Fig. S9). To determine if YB-1 is activated in response to injury, we performed optic nerve crushes and collected the ipsilateral and contralateral retinas 48 hours later. We observed baseline phosphorylation of YB-1 in the contralateral retina, and this was visibly increased in response to optic nerve crush (Fig. 5A). We also found that histones H3 and H4, as well as mixed histones, could induce phosphorylation of YB-1 (Fig. 5B, quantification of data is in Fig. S10). We also noted that YB-1 expression had a direct impact on neurite outgrowth, as overexpression of YB-1 in cortical neurons resulted in shorter neurites compared to controls (Fig. 5C). As the application of histones to distal neurites is sufficient to inhibit neurite outgrowth (Figs. 3H), and injuring the optic nerve will subsequently lead to elevation of pYB-1 in the retinal ganglion cell layer, it appears that histones may induce phosphorylation of YB-1 through retrograde signaling. To test this hypothesis, the neurite compartments of microfluidic chambers were treated with histones coupled to 6 µm silicone beads, which prevents their diffusion into the somal compartment, while preserving their ability to interact with the neurites. When the histone-conjugated beads were applied to the neurites, we observed increased pYB-1 levels in the neuronal cell bodies, confirming a retrograde signal (Fig. 5D). Interestingly, the neurites treated with the histone-conjugated beads displayed prominent varicosities (Fig. 5E4, while neurites treated with unconjugated beads were unaffected (Fig. 5E3.). This not only demonstrates that the beads had no toxic effects, but also provides further evidence that histones negatively affect axonal function and integrity, as these varicosities are indicative of disruptions in axonal transport and impending degeneration. These results demonstrate that CNS injury can induce YB-1 phosphorylation in neurons and given our data showing that histone levels are elevated following optic nerve crush (Fig. 5A), it is highly probable that this occurs through histone-mediated retrograde signaling. The induction of neurite degeneration by the histone-coupled beads suggests that the histones are binding to a receptor. The Nogo receptor (NgR) has been shown to bind the 3 major myelin-associated inhibitors, leading to inhibition of neurite outgrowth and growth cone collapse (20-23). Using microfluidic chambers, we treated the somal compartments with NgR siRNA, waited 48 hours and then added histones to the neurite compartment and waited another 48 hours. We found that siRNA knockdown of NgR provided no protection from the effects of histones, as histone-mediated inhibition was comparable to that seen in a scrambled siRNA control (SC) with histone treatment (Fig. 5F). We then tested for other potential receptors. Rodent cortical neurons have been shown to express Toll-like receptor 2 and 4 (TLR2 and 4; 11). We treated cortical neurons with siRNA for TLR2 and TLR4, either individually or in combination. When the neurite compartment was treated with histones following the knockdown of TLR2, but not TLR4, we observed a partial block of the inhibitory effects of histones (see Fig. 5F&G). To confirm that our siRNA were working, we performed RT-PCR and found that after 48hrs, siRNA for TLR2 reduced TLR2 mRNA levels by nearly 40% compared to SC siRNA-treated cortical neurons. This reduction in TLR2 mRNA persisted for up to 96hrs and in some cases increased to nearly 65% reduction, which covers the time frame of our experiment. For TLR4, the siRNA was not as potent resulting in only 20% reduction in mRNA levels compared to SC siRNA-treated cortical neurons after 96hrs, with no significant decrease in TLR4 mRNA detected at 48hrs. This difference in siRNA performance could account for why TLR2, but not TLR4, is having an effect in our neurite outgrowth assays with histones.

**Legend 5.**
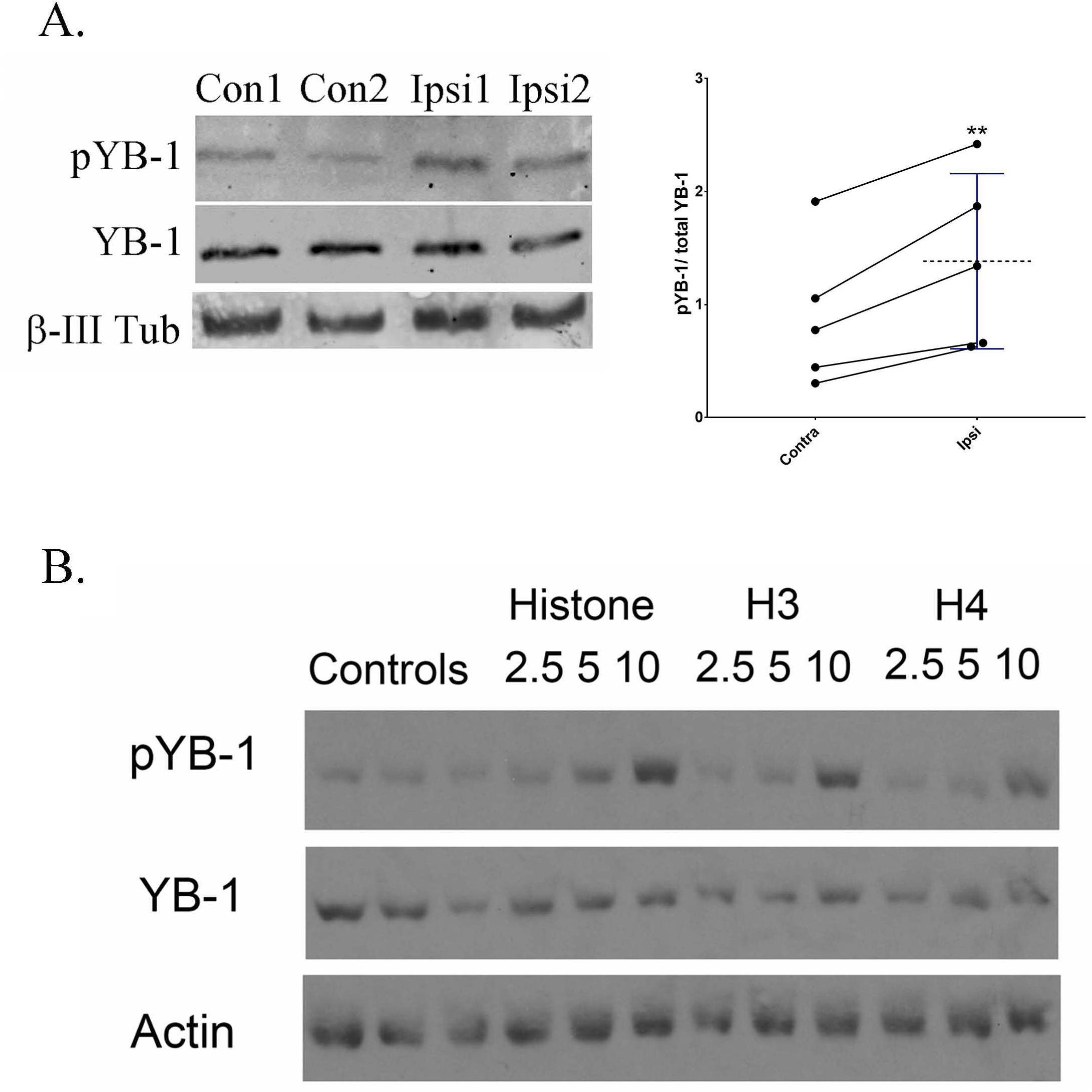
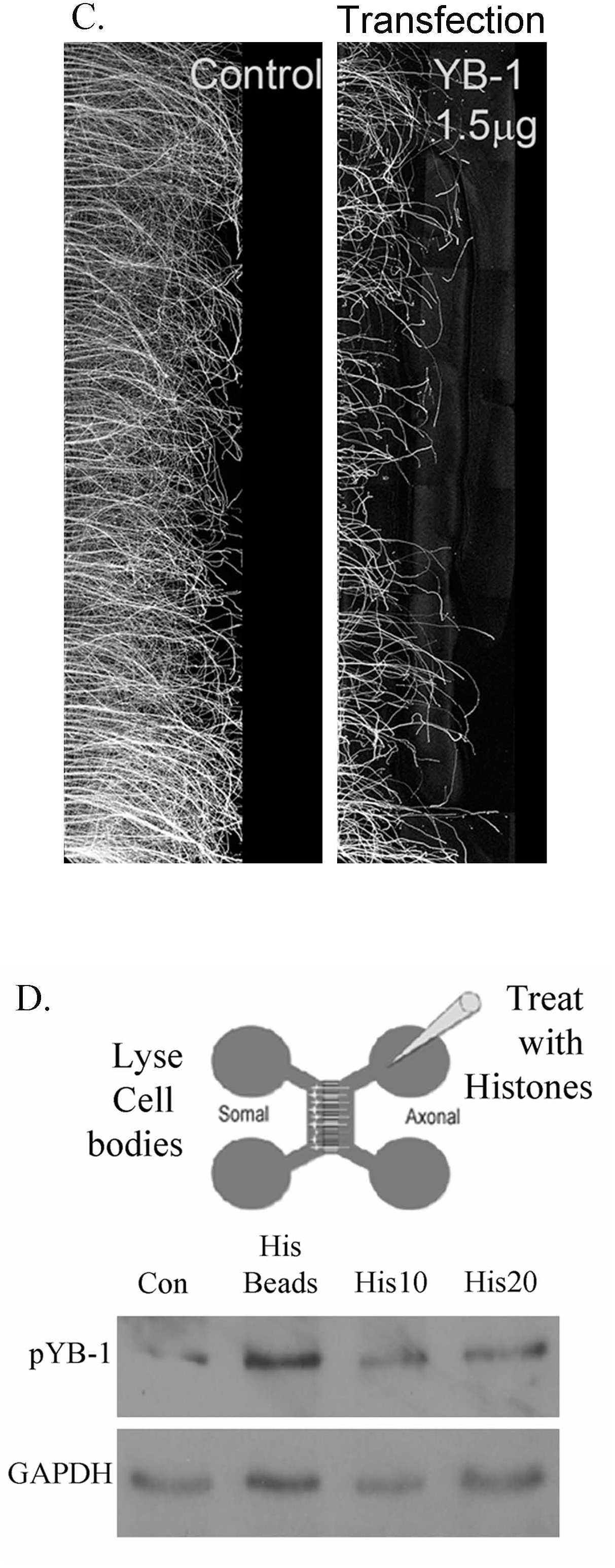
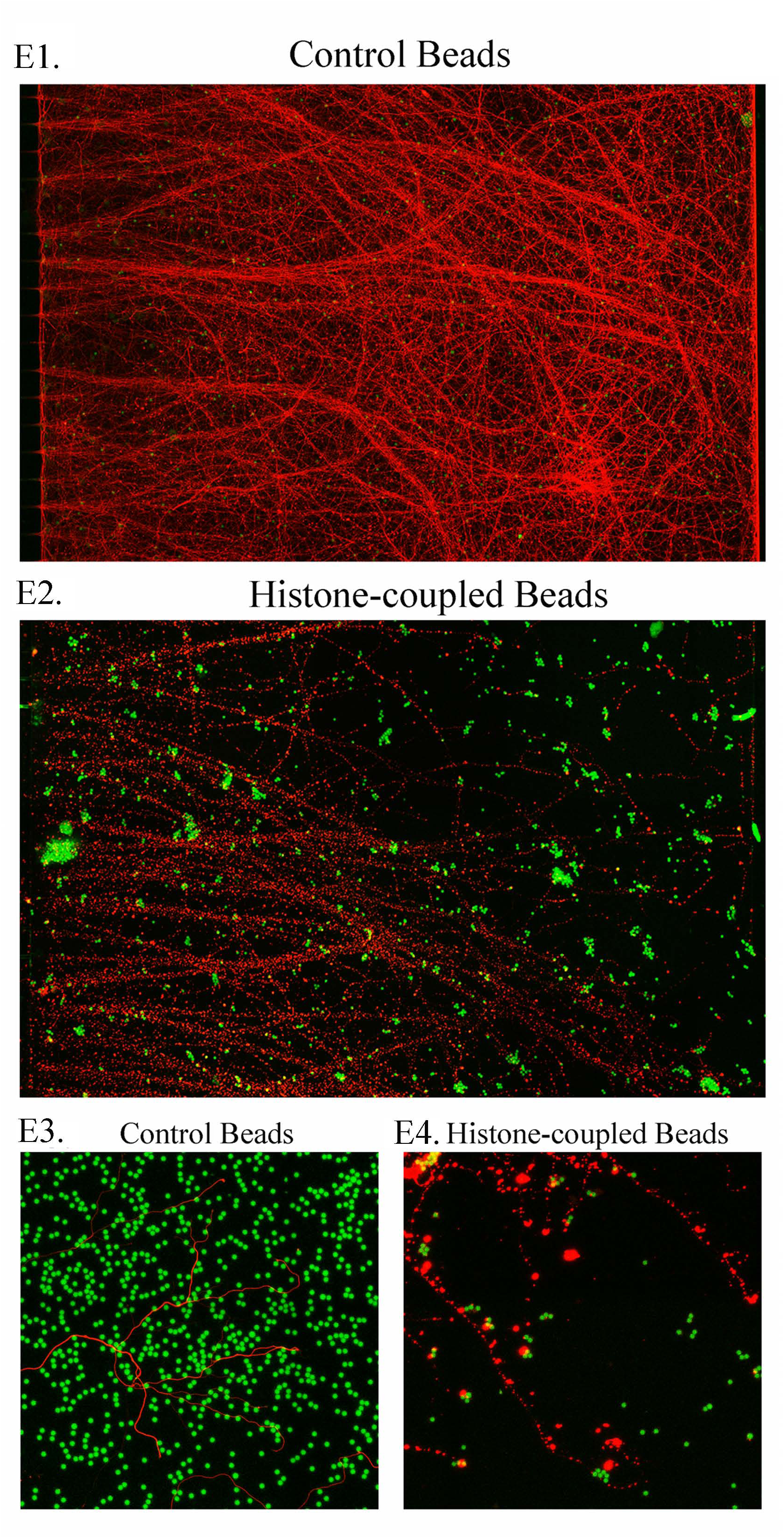
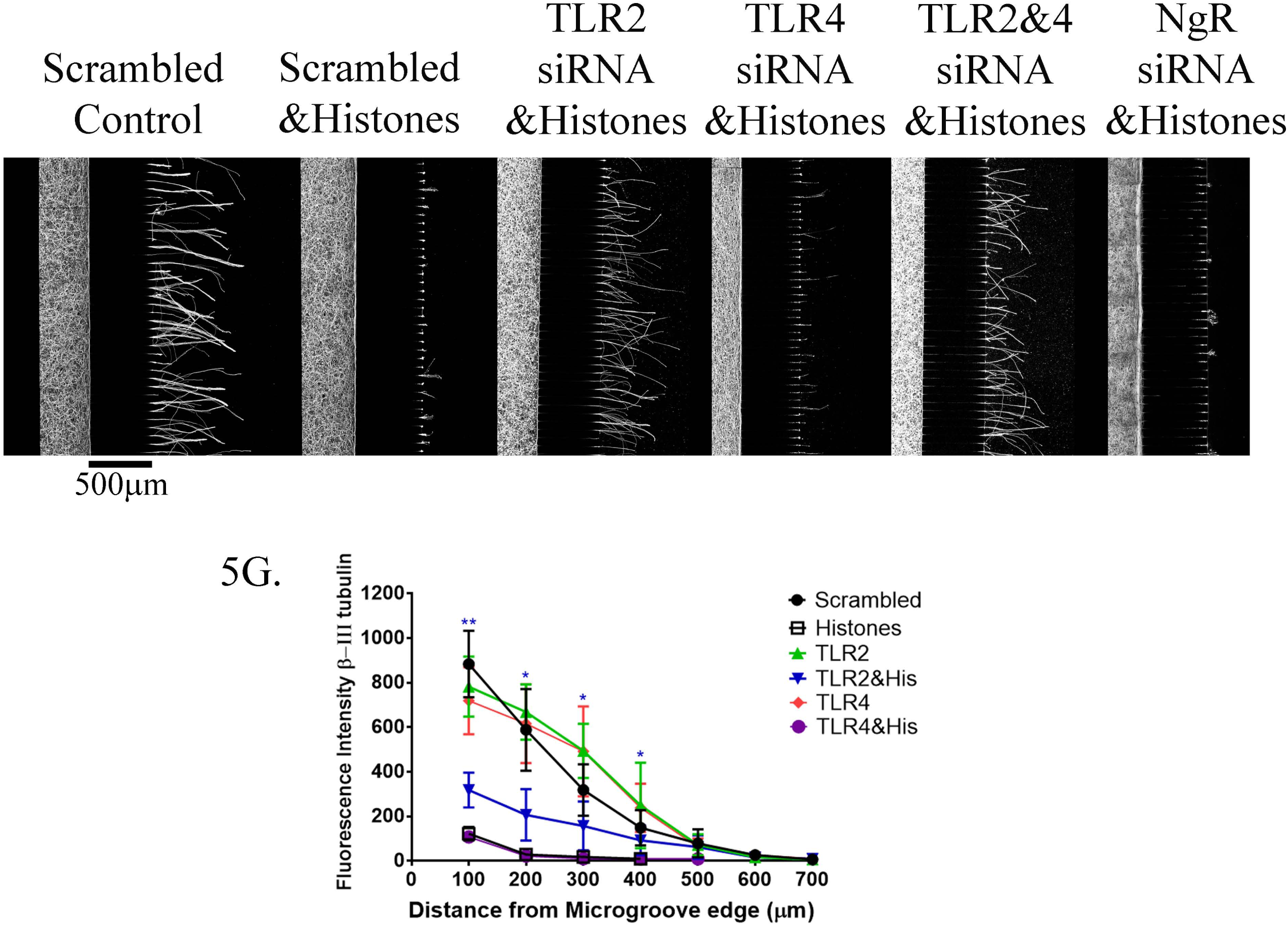
Histones inhibit neurite outgrowth by an YB-1 mediated mechanism and Toll-like receptor (TLR) 2 knockdown by siRNA provides protection from the inhibitory effect of histones. **A.** Cortical neurons treated with mixed Histones, H3 or H4 (2.5,5, or 10µg/ml) induce elevation in phosphorylated YB-1 (pYB-1), normalized to YB-1 and Actin levels. **B.** To confirm if pYB-1 are elevated after injury inducing histone release in the ONC, we crushed the axons of the optic nerve and 48hrs later collected the retinal cell layer, ipsilateral side (Ipsi 1&2), and collected the retinal layer of the non-injured, contralateral side (Con1&2) in the same animal and prepared them for western blots. We found significantly elevated levels of pYB-1 normalized to total YB-1 on the injured (Ipsi) side compared to the non-injured (Contra) side. Statistics is paired t-test, **p<0.01. **C.** Overexpressing the YB-1 plasmid in cortical neurons by NEON transfection and plating in microfluidic chambers results in shorter neurite outgrowth in a dose dependent manner as determined by β-III-tubulin staining. **D.** Cartoon of a microfluidic chamber diagrammatically showing where we apply the histones or to the neurites and then lysed the cell bodies 48 hours later, on the opposing side. Cortical neurons plated in microfluidic chambers for 4-5 days grow long neurites across the microgroove. Using histones or fluorescent microbeads that are covalently coupled to histones (or non-coupled for controls) and too large to cross the microgrooves, or with 10 or 20µg/ml mixed Histones (His10 or His20) are applied to the neurite compartment (axonal) only and we wait 48hours before lysing the cell body compartment (somal) with 2XRIPA buffer and run a western blot. We detect elevation of pYB-1 in the cell bodies normalized to GAPDH. **E1.** Non-coupled beads (green) had no effect on neurite outgrowth (β-III-tubulin in red), **E3** shows higher magnification. **E2**. Histone-coupled beads (green) appear more aggregated compared to non-coupled beads and significantly inhibited neurite outgrowth. **E4.** higher magnification of the beaded BIII-tubulin (red) with histone-coupled beads. **F.** Neurite outgrowth in chambers using siRNA applied to cell bodies compartment (Left of the microgrooves). Imaged at 20X, micrometer of 500μm bottom of Scrambled control. Controls were treated with scrambled siRNA, 48hrs after siRNA application 20μg/ml histones were applied to neurite chamber (Right of the microgroove) and incubated for 24hrs. Neurites were immunostained with β-III-tubulin and we quantified the neurite length by ImageJ analysis. **G.** Graphs of the average quantified length are shown. Statisitcs are t-tests, where we compared scrambled with Histones to TLR2 siRNA with Histones, and we see a significant difference, *p<0.05 and **p<0.01.

To determine if APC could promote axonal regeneration *in vivo*, APC-containing gelfoam was applied to the injury site following ONC in rats (Figs. 6A&B). Retinal ganglion cell axons were anterogradely labeled with cholera toxin B-subunit (CTB) at 14 days after injury, and optic nerves were chemically cleared using the 3-DISCO technique (15). Crush sites were identified using a novel technique, the second harmonic generation, which allows visualization of collagen fibrils (Figs. 6E&E’, F&F’) (16&17), and axonal regeneration was quantified by fluorescent intensity from the crush site at 100 µm increments (Fig. 6C) and volumetric analysis of CTB-labeled axons (Figs. 6D & S11). Control animals treated with saline-containing gelfoam displayed limited growth and short CTB-labeled axons crossing the lesion site (Fig. 6A). In APC-treated animals, however, we observed a significant increase in axonal regeneration beyond the site of injury (Figs. 6B & another representative figure with 3-D projection of fibers in S12).

**Legend 6.**
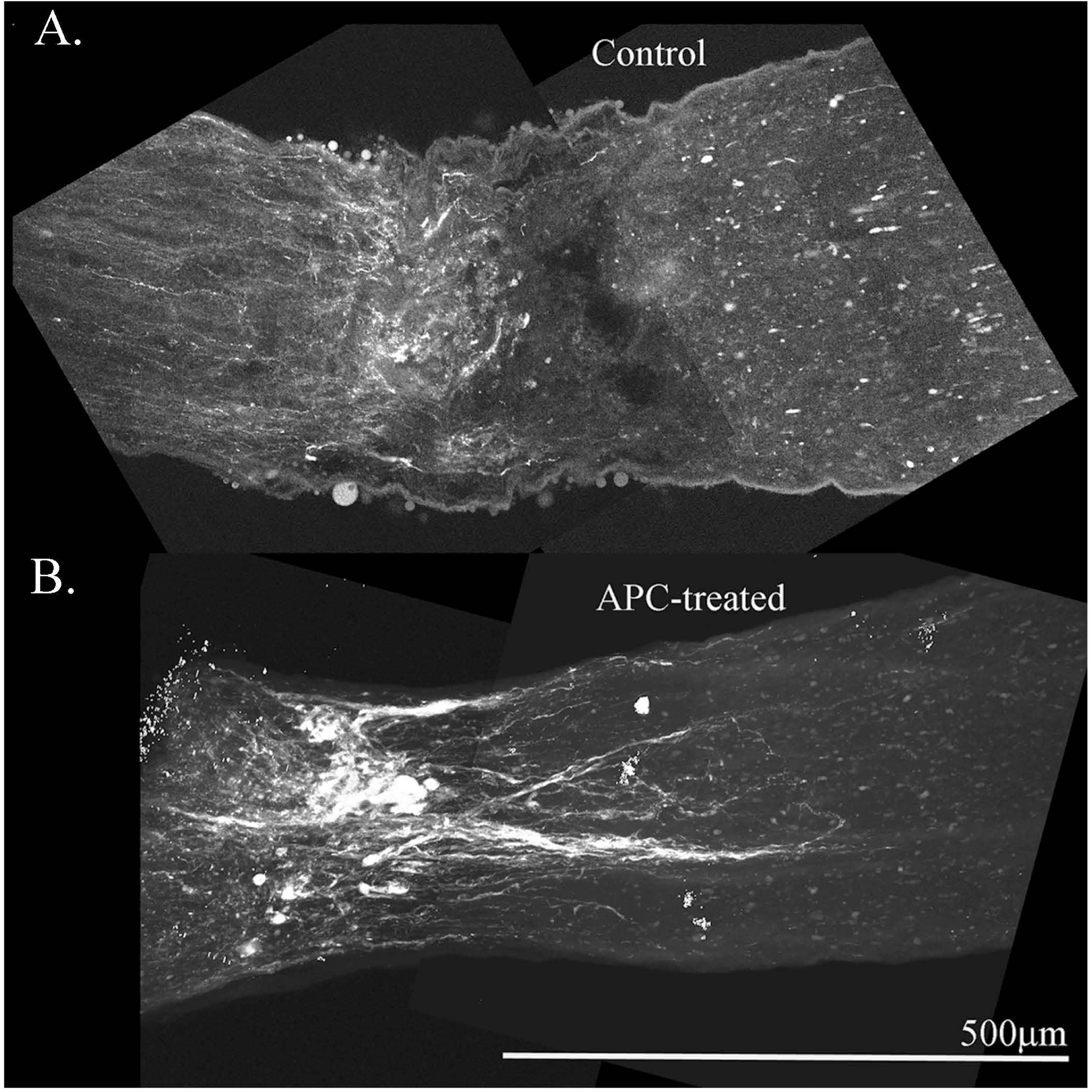
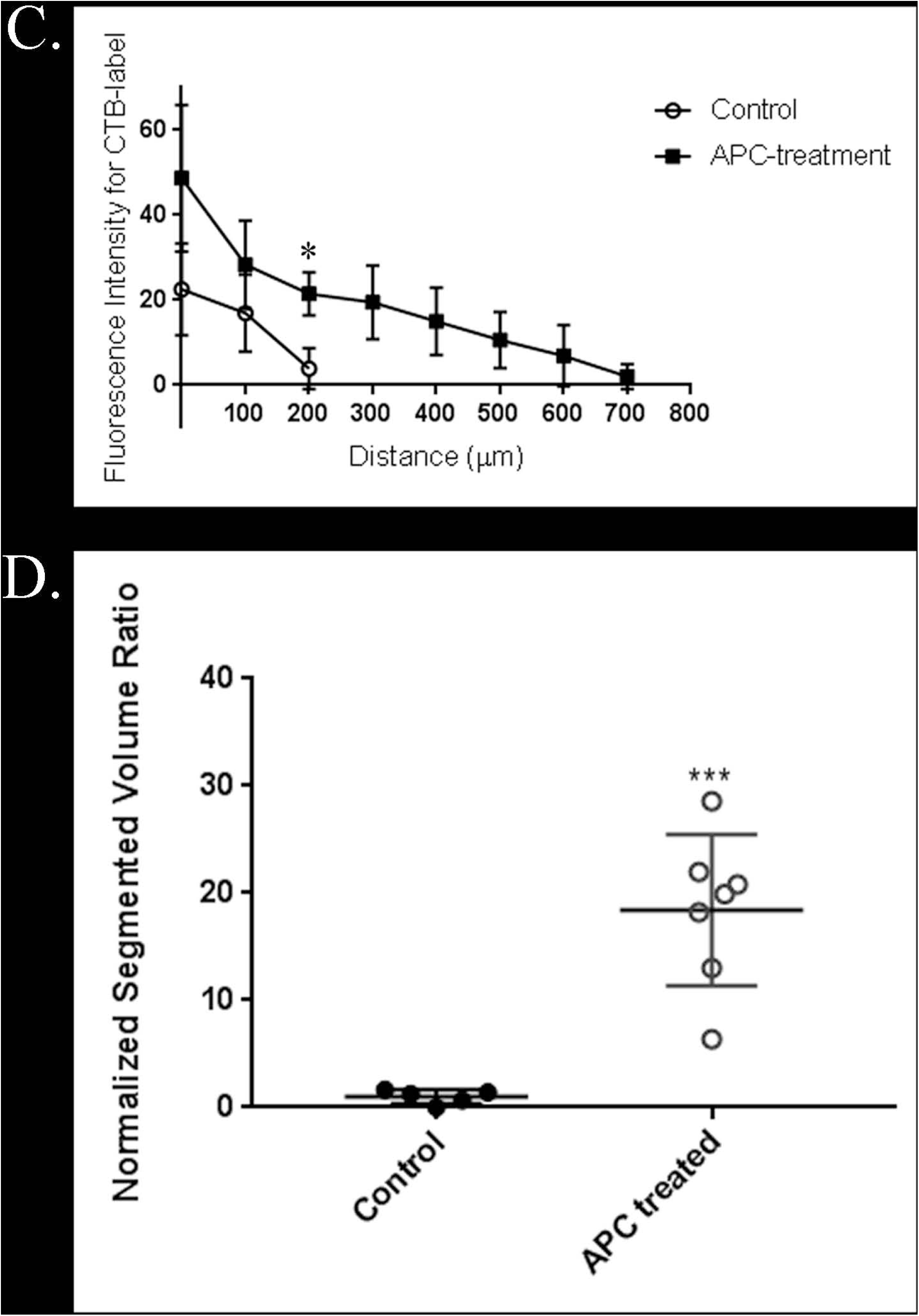
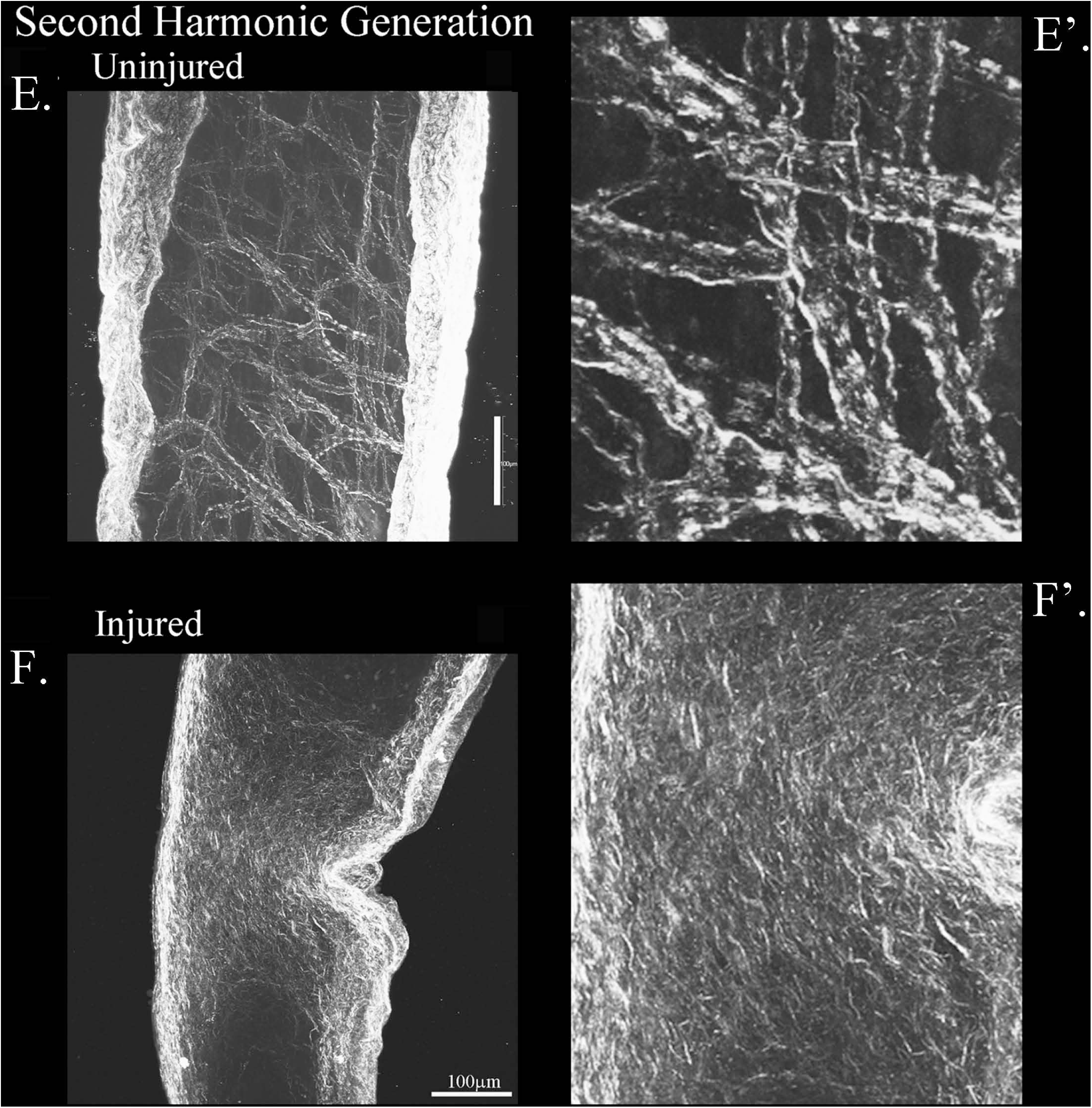
**A.** Using the rat ONC model for axonal regeneration in the CNS, crushed axons without treatment (Control) have CTB labeled axons abruptly halt at the lesion site in this chemically cleared whole nerve. **B.** Representative optic nerve treated with APC soaked in gelfoam immediately after crushing the axons, the APC-gelfoam is placed over the injury site. In this chemically cleared nerve CTB labeled axons are observed growing through the lesion site. Panel A & B are approximately 540 optical slices at 0.45µm increments that is merged together into a two-dimensional represented maximum projection intensity image. Imaged at 25X on an Olympus Multiphoton microscope and scale bar of 500µm is for Figures A & B. **C.**Using Image J we quantitated the axonal regeneration for Control (n=5, white circle) and APC-treated (n=7, black square) every 100microns from the edge of the crush site. There is a significant difference in the extent of regeneration at 200microns, and no regenerating fibers were detected past that point in Control animals. **D.** Using Amira 3D maximum intensity projection of the nerve showing crush site and regenerating neurites labelled with CTB, we determined the Normalized Segmented Volume ratio. Line plot shows ratio of segmented neurite area over total nerve area per cross section along the nerve and we see a significant difference (***, p<0.001) with APC-treatment. Each cross section slice is .497 micrometers. **E.** Using Second Harmonic Generation imaging on Multi-photon microscope, collagen fibers auto-fluoresce. In uninjured whole chemically cleared nerves, we can see the collagen bundles, which the fibers are shown in enhanced detail in Figure **E’**. **F.** Injured optic nerves display crushed collagen fibers at the injury site, helping to elucidate the injury site and it is shown in an enhanced region in Fig. **F’**. Scale bar is 100µm in Figures E. and F.

## Discussion

Histones lie at the very core of chromatin structure and function, but here we show that they have a very different role outside the nucleus. Histones are released into the extracellular environment following CNS injury, and they act as potent inhibitors of axonal regeneration both *in vitro* and *in vivo*. Extracellular histones produce a degree of inhibition that is comparable to that elicited by myelin-associated inhibitors and CSPGs (22,24,25), and we therefore propose that histones should be considered a third major class of inhibitory molecules within the CNS. Eliminating inhibitory factors from the CNS environment has been widely used as a strategy to enhance axonal regeneration, but the impact has often been minimal. Triple mutant mice lacking expression of myelin-associated glycoprotein (MAG), Nogo, and oligodendrocyte myelin glycoprotein did not exhibit spontaneous regeneration of corticospinal and raphespinal axons after spinal cord injury (5). Similarly, elimination of CSPGs through treatment with chondroitinase ABC (chABC) resulted in only modest regeneration of dorsal column and corticospinal axons (maximum of 4 mm) following dorsal column lesions (26). The presence and persistence of extracellular histones at the site of injury could therefore account for the limited axonal regeneration that was observed in these studies.

Having established that extracellular histones inhibit axonal regeneration, we studied their mechanism of action, and our observation that APC can block the effects of histones provides some initial insight. Since APC cleaves histones, we propose that histone proteins are directly responsible for mediating inhibition. This is further supported by our observation that APC cannot overcome inhibition by MAG in a neurite outgrowth assay (Fig. S13), which demonstrates that APC does not enhance growth through non-specific degradation of myelin-mediated inhibitors or effects on the neurons themselves. Our results show that histones induce phosphorylation of YB1 through retrograde signaling, which suggests that this effect is receptor-mediated. Receptor ablation experiments show that TLR2, but not NgR or TLR4 mediate the observed effects. Histones have been shown to activate intracellular signaling by binding to cell-surface receptors such as TLR2 and TLR4 in other systems (12). It has also been reported that YB-1 can promote microtubule assembly (27), and so it is possible that histones may activate RhoA through phosphorylation of YB-1.

The addition of histones to the list of inhibitory molecules in the CNS reinforces the prevailing view that combinatorial therapies will be necessary to promote axonal regeneration and functional recovery following spinal cord injury. Elevation of intracellular cyclic AMP can overcome inhibition by myelin-associated inhibitors and CSPGs, and we have found that it can reverse inhibition by histones as well (see Fig. S1C) (28, 29), which suggests that rolipram or other phosphodiesterase inhibitors could be an effective way to target all three classes of inhibitors. Histones could also be neutralized using APC, which would digest the histones within the lesion site and create a more permissive environment for regeneration. It should be noted that this treatment would not affect histones in intact cells, which means that APC could be selective, with few adverse effects *in vivo*. This approach could be particularly effective when used in combination with chABC and other interventions that modify the extracellular environment, such as cell transplantation. The findings presented in this study highlight the complexity underlying the challenge of promoting axonal regeneration in the CNS, but they also reveal new opportunities for developing mechanism-based treatments for spinal cord injury relevant to humans.

## Supporting information

Full Supplement

## Acknowledgements

This work was supported by funding by the Specialized Neuroscience Research Program (3U54NS041073, NINDS-NIH) and R01NS037060 (NINDS-NIH) awarded to M.T. Filbin, Research Centers in Minority Institutions Programs (RR00307, NCRR-NIH) awarded to Hunter College, R01GM054508 and R01GM137056, and Systems Biology Center grant P50GM071558 awarded to R. Iyengar. F32NS054511 awarded to M.M. Siddiq. NYS Spinal Cord Injury Research Program Contract number #C34460GG awarded to RI and MMS. The authors would like to thank Drs. Robert Blitzer, Joseph Goldfarb and Wilfredo Mellado for critically reviewing this paper, and Gomathi Jayaraman and Som Prabha for their excellent technical assistance. Multiphoton microscopy was performed in the Microscopy CORE at the Icahn School of Medicine at Mount Sinai and was supported with funding from NIH Shared Instrumentation Grant 1S10RR026639-01. The authors gratefully acknowledge the International Spinal cord Injury Biobank (ISCIB) for generously providing the human specimens used in this project. *This paper is dedicated to the memory of Marie T. Filbin*.

## Author Contributions

MMS performed all experiments for this project and did all the microscopic work.

SSH assisted with data analysis and validation for reproducibility of experiments.

YZ performed and analyzed the Panomics Assay, and assisted with experiments.

EN assisted with surgeries and western blots, and validated experimental results.

VR, ELR and RET assisted with surgeries and all in vitro experiments.

JH performed CSF collection from mice.

RH performed the volumetric analysis on blinded samples of ONC.

JH analyzed and performed all computational work for the bulk-mRNA-sequencing data.

BKK collected human CSF samples.

AV and BKK curated and maintained all human CSF samples.

SET assisted with analysis of CSF samples and critical review of the manuscript.

IM provided recombinant histones and critical review of the manuscript.

RS performed the bulk mRNA-seq.

MMS, RI and MTF conceptualized the project and provided necessary funding acquisition.

MMS, SSH and RI wrote and edited the manuscript.

## Supplementary

**Figure S1.**
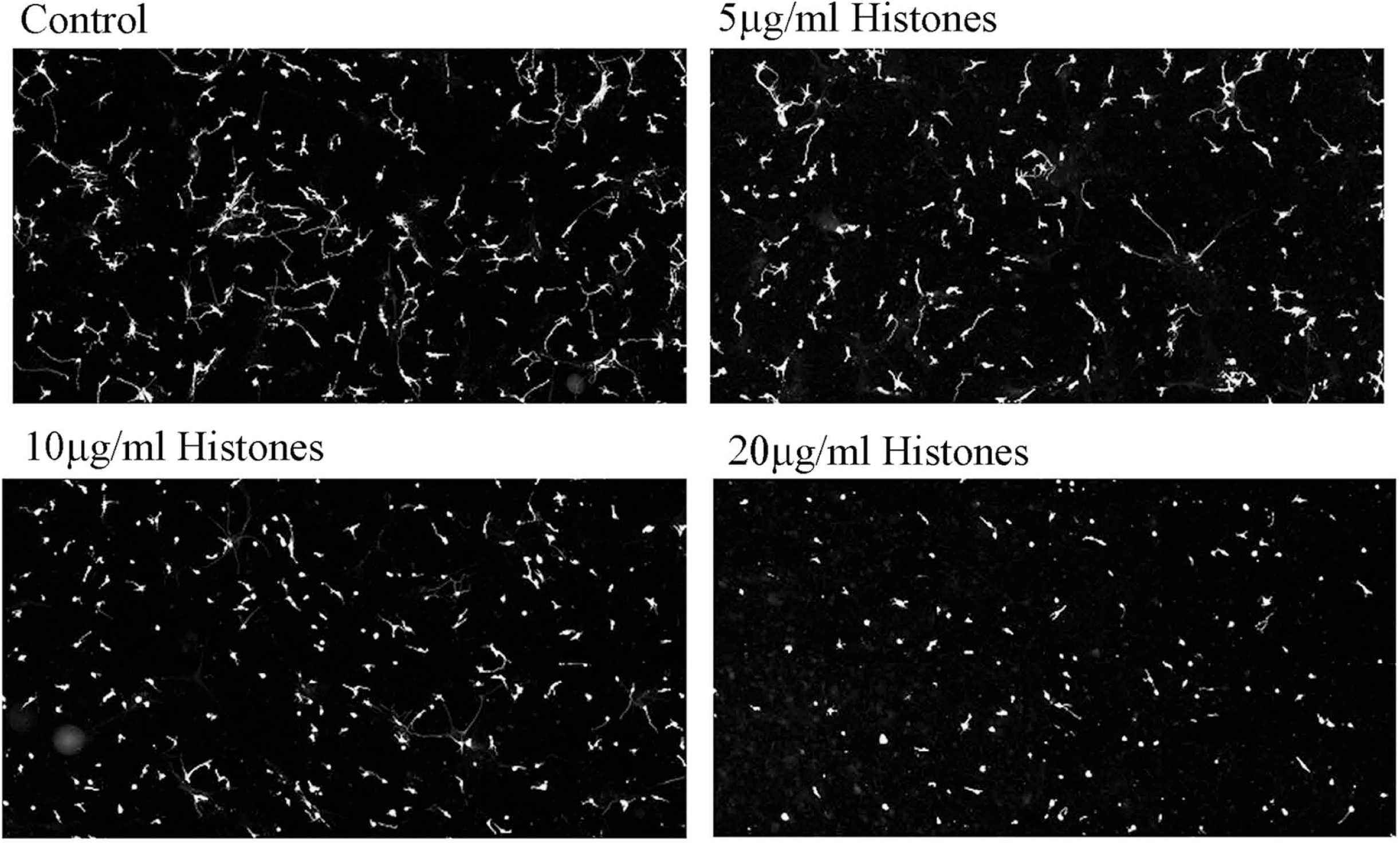
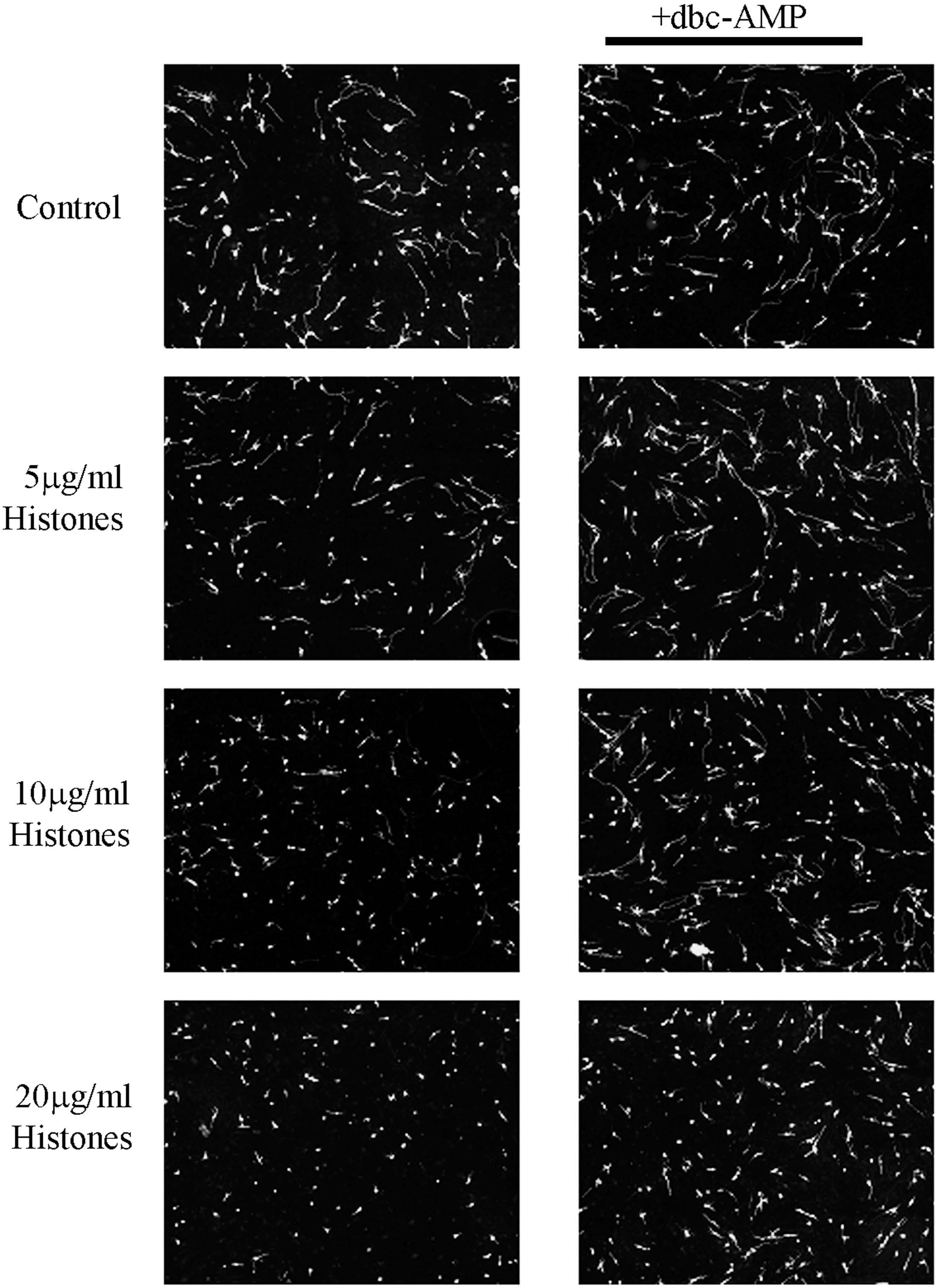
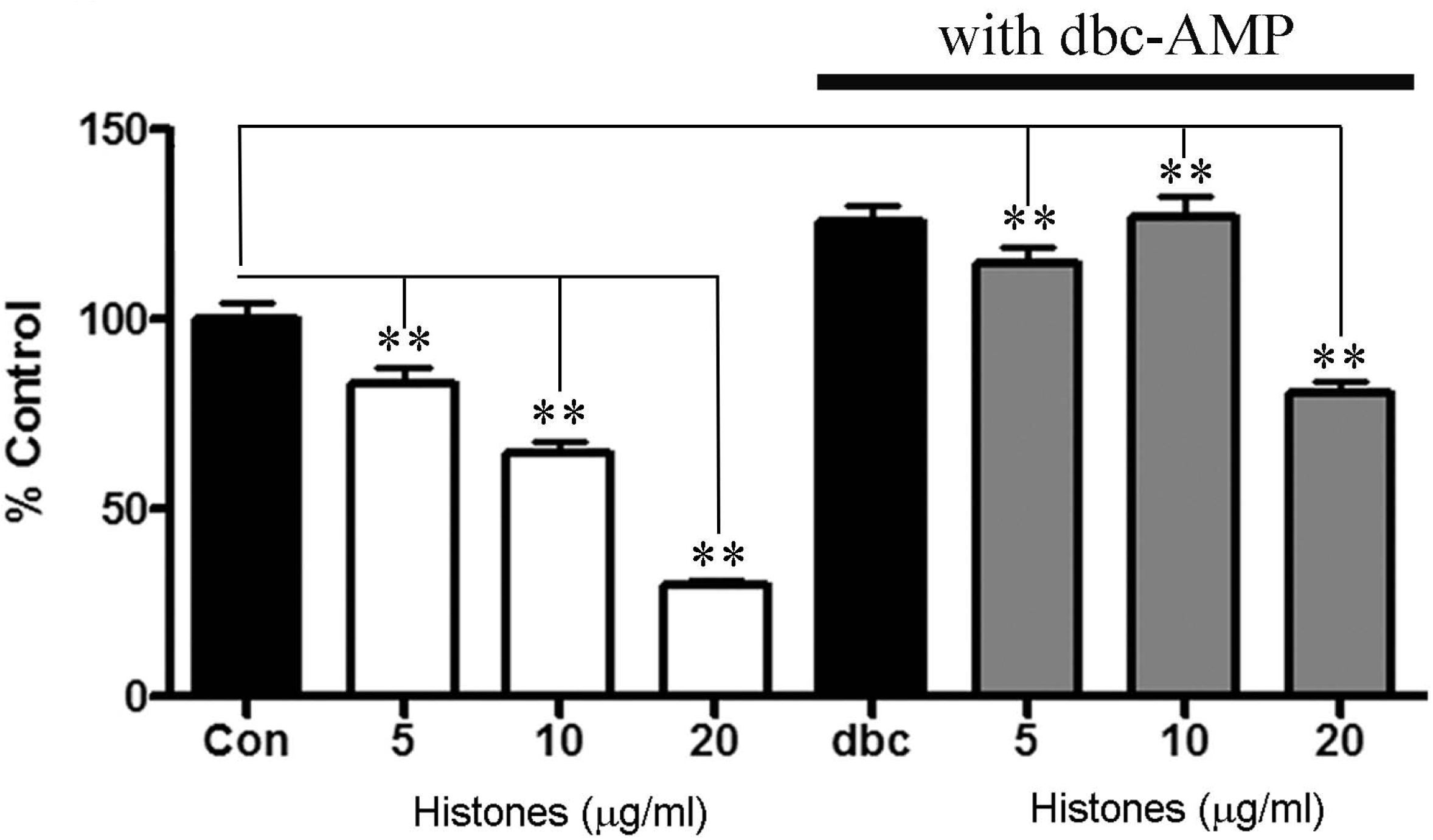
Extracellular histones inhibit neurite outgrowth on a permissive substrate CHO monolayers. A. Representative images showing the whole field imaged and quantitated. These are cortical neurons on CHO monolayers stained with β-III tubulin and a dose-dependent inhibition of neurite outgrowth by mixed population of histones from calf thymus (5-20µg/ml) was observed. B. Representative images of cortical neurons on CHO monolayers stained with β-III tubulin and a dose-dependent inhibition of neurite outgrowth by mixed population of histones from calf thymus (5-20µg/ml) and histones applied with 1mM dbc-AMP, where dbc- AMP can overcome the inhibitory effect to neurite outgrowth. C. Quantification for the images in Figure S1B, each bar is the average of there independent experiments. For statistics we used ANOVA and compared all treatments to control, no treatment, **p<0.01.

**Figure S2.**
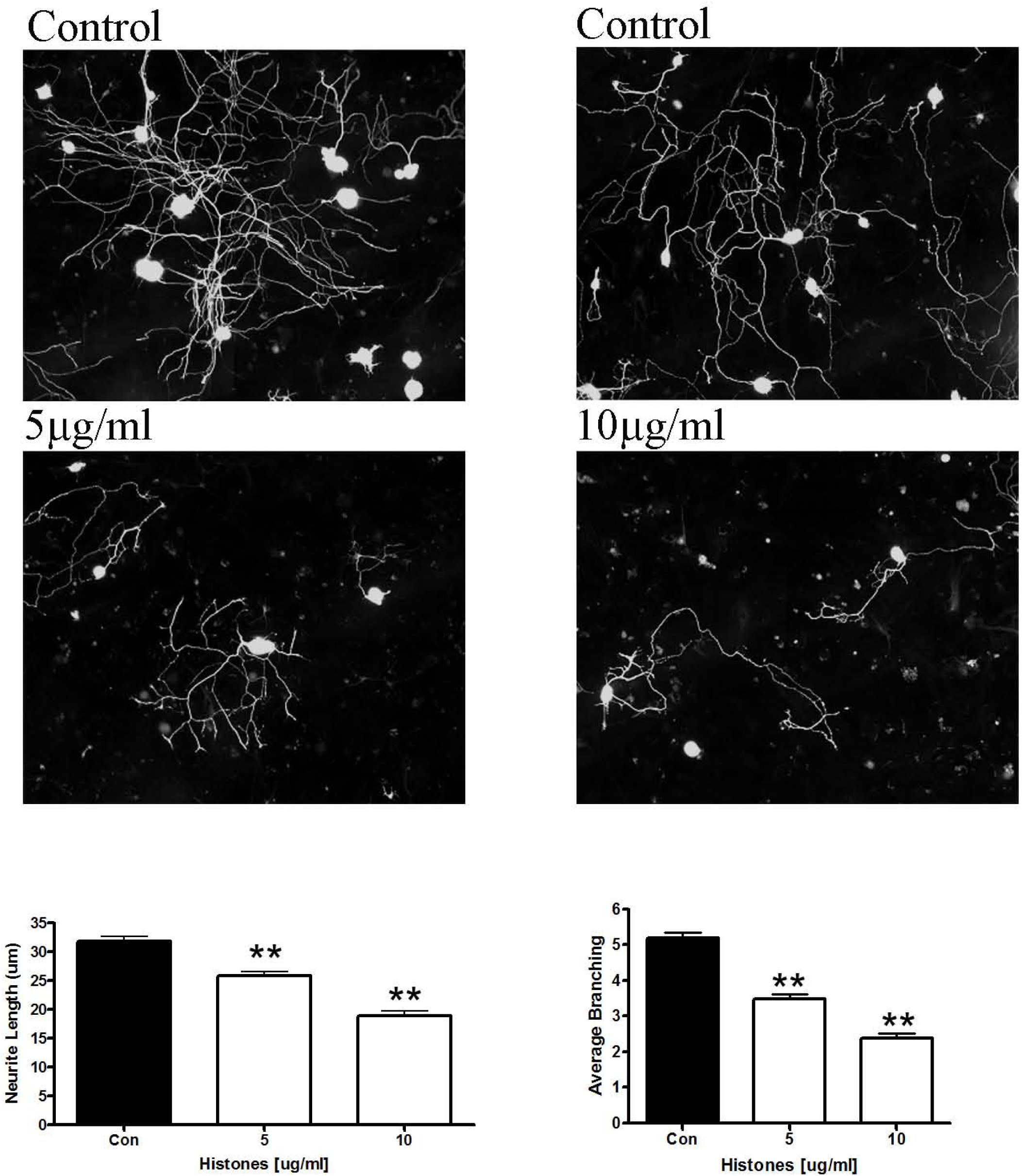
Extracellular histones are inhibitory to neurite outgrowth and braching in primary dorsal root ganglion. Dorsal root ganglion (DRG) neurons grow shorter neurites and have fewer branches in the presence of extracellular histones. P5 rat DRG neurons are plated on a permissive layer of CHO monolayers with or without extracellular histones. Histone-treatment results in shorter neurites and we also quantified the average number of branches which was significantly reduced in the presence of histones. All statistics are performed by ANOVA, **p<0.01.

**Figure S3.**
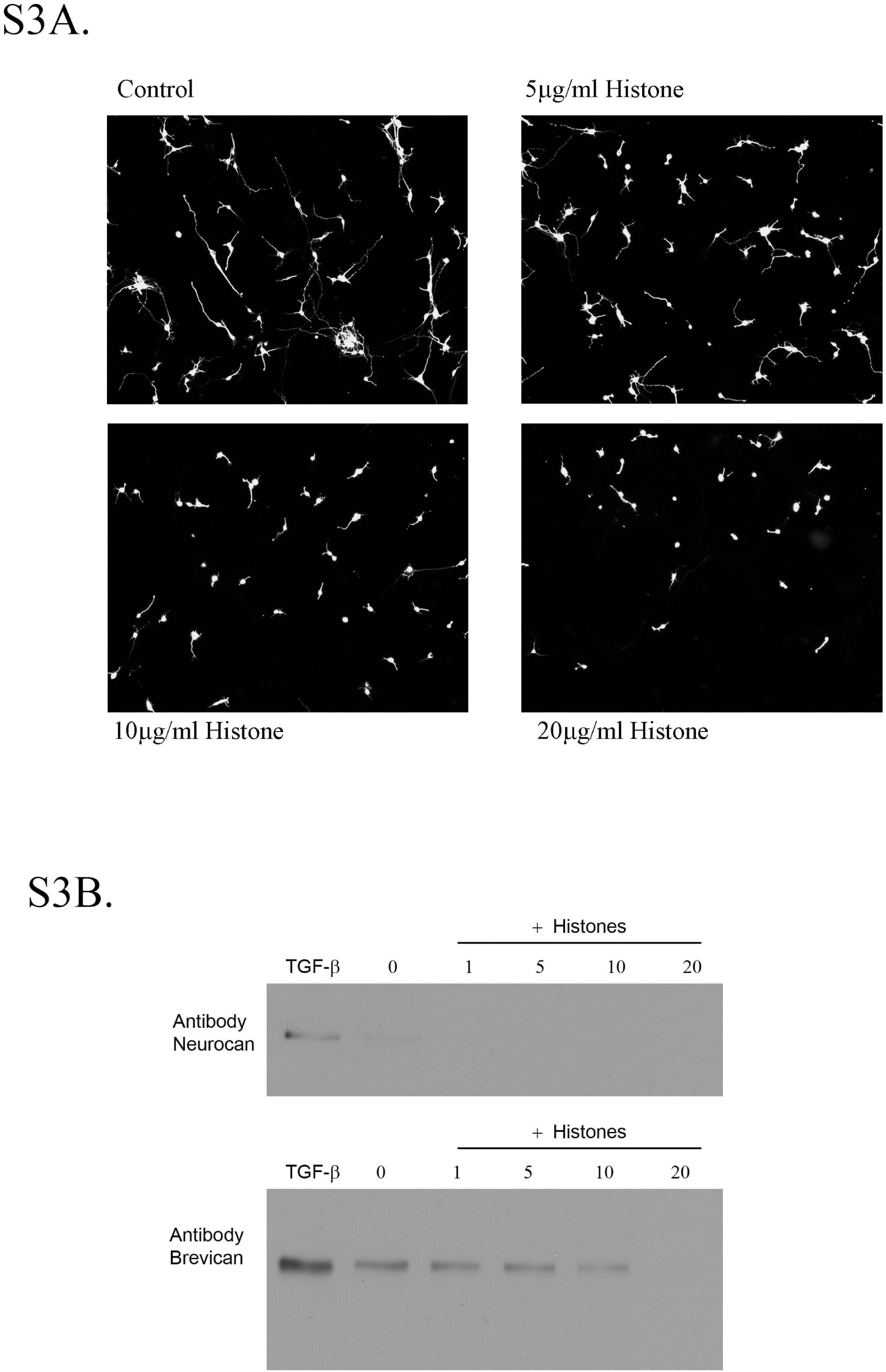
Extracellular histones inhibit neurite outgrowth on a permissive GFAP+ Astrocytic monolayer and do not affect CSPG secretion. A. Primary rat astrocytes grow on PLL-coated glass 8 wells microscope slides to confluency. We applied histones in the concentrations indicated and added cortical neurons at the same time, we incubated for 24hrs, stained for β-III-tubulin and observed a dose-dependent inhibition of neurite outgrowth. B. Astrocyte cultures were treated with either mixed histone population or with TGF-β, which is known to increase secretion of CSPGs. Compared to TGF-β, histones did not up-regulate secretion of either neurocan or brevican.

**Figure S4.**
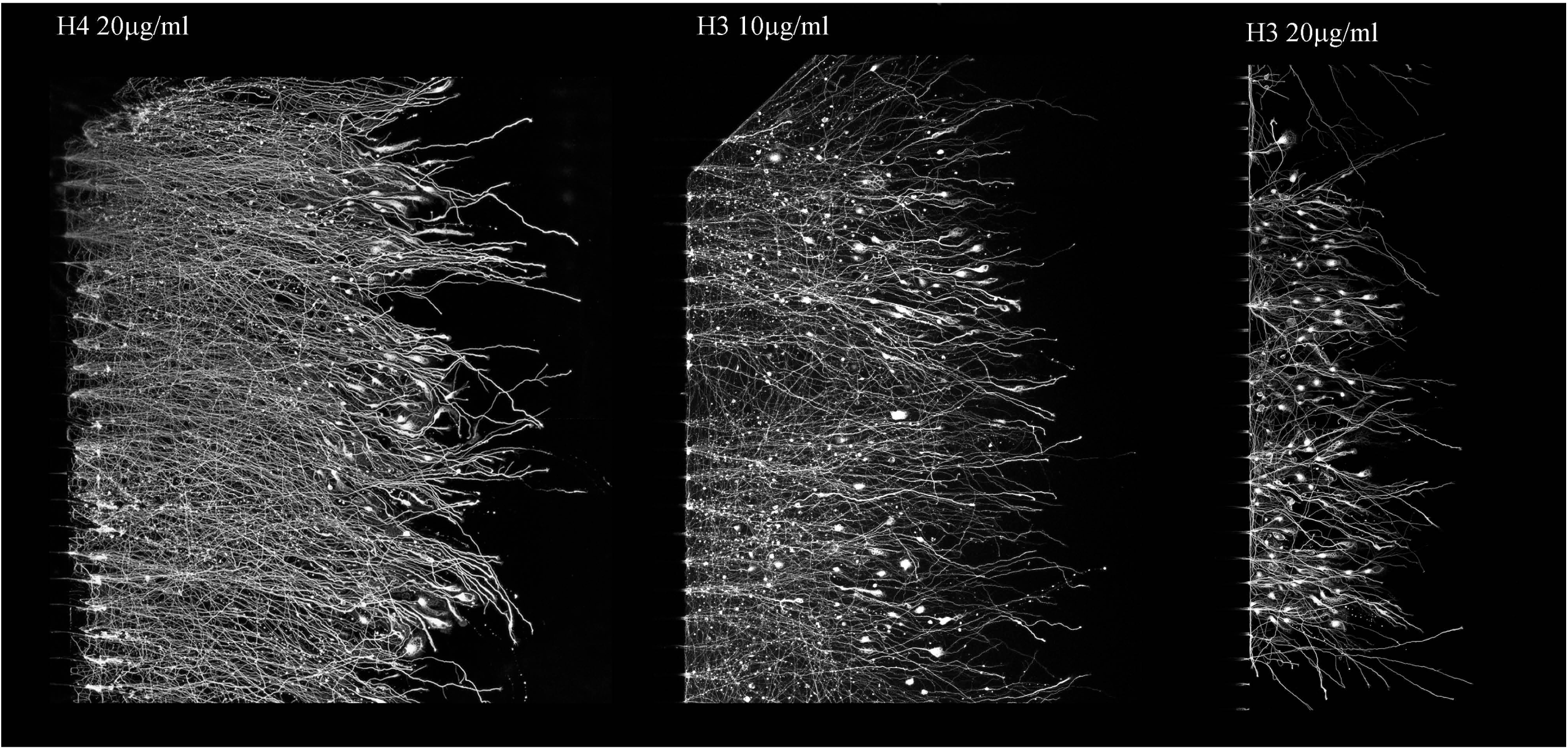
Recombinant H3 and H4 histone isoforms are inhibitory to neurite outgrowth and contribute to dystrophic bulb formation. Using microfluidic chambers with cortical neurons treated with either recombinant H3 or H4 isoforms, we observed that H3 had a more robust effect on inhibiting neurite outgrowth and promoted dystrophic bulb formation. Though H4 also resulted in slightly shorter neurites and promoted dystrophic bulb formation.

**Figure S5.**
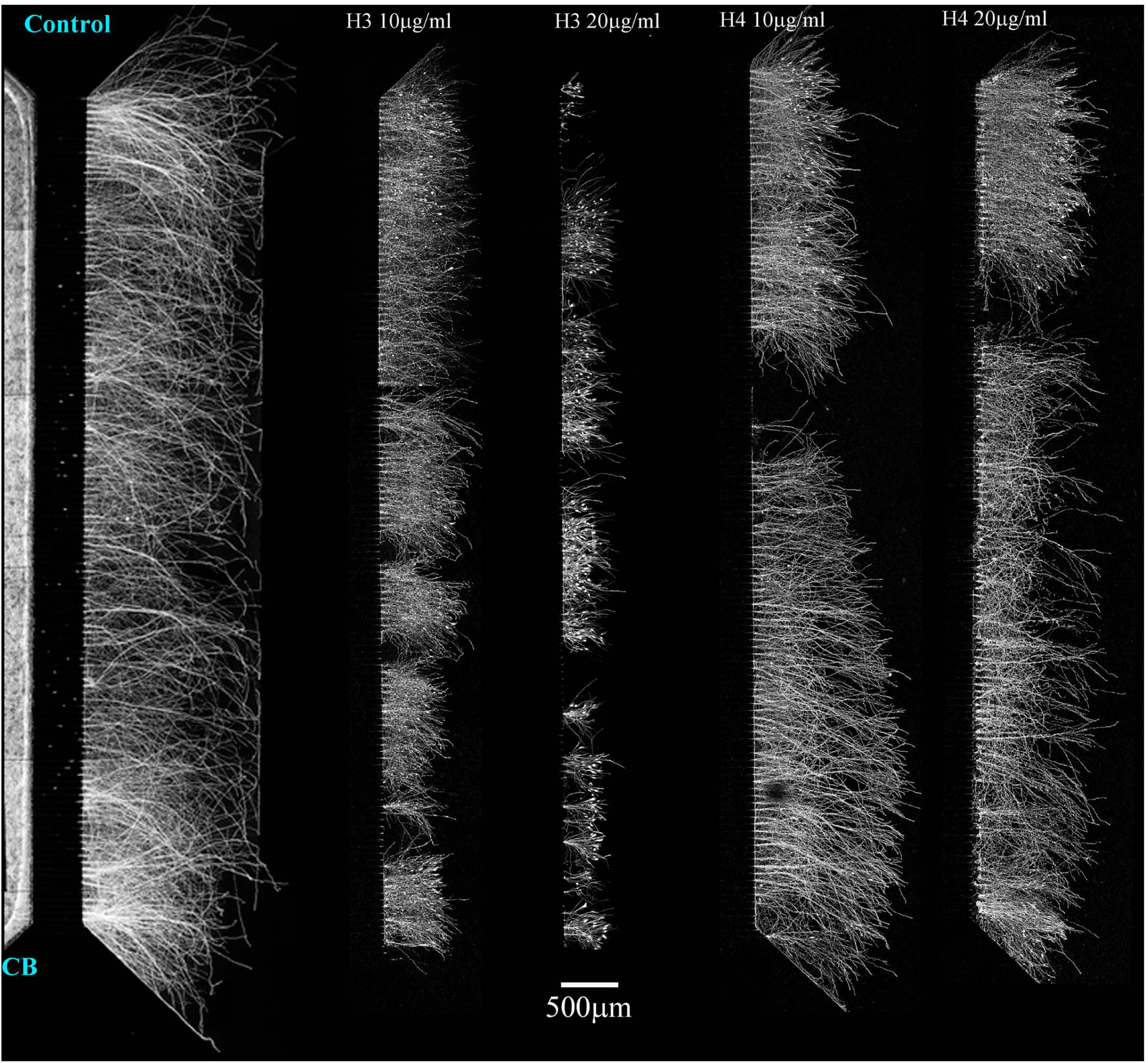
Extracellular histones are inhibitory to neurite outgrowth on PLL-coated microfluidic chambers. Using PLL-coated microfluidic chambers, cortical neurons cell bodies (CB) grow long neurites across the 450µM microgroove as detected by β-III tubulin. Treating the neurite growing compartment with either recombinant H3 or H4 Histones resulted in significantly shorter neurites, with H3 having a more potent effect. Representative images showing the full chamber.

**Figure S6.**
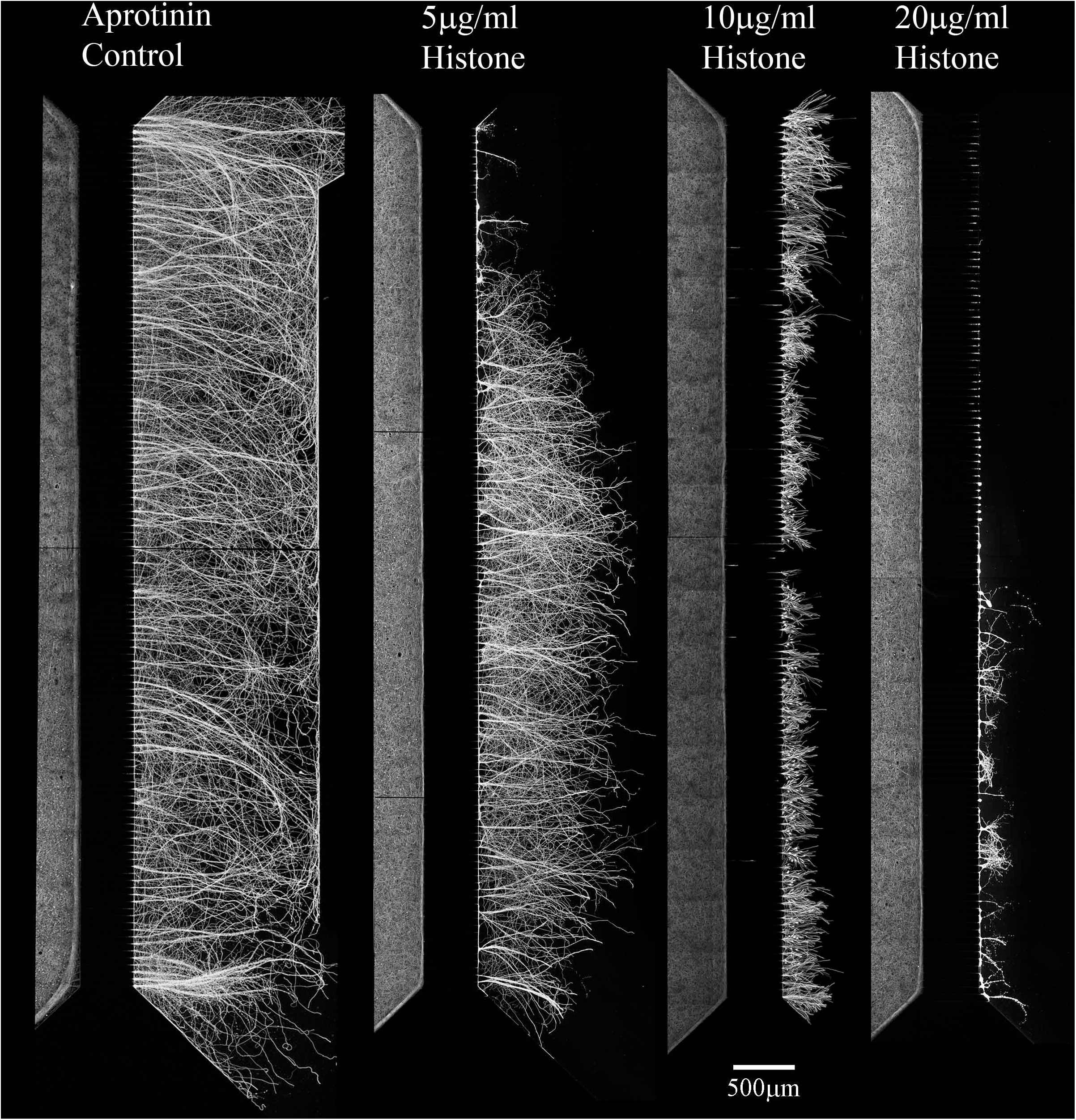
Mixed population of histone are inhibitory to neurite outgrowth in a dose-dependent fashion. Cortical neurons plated on PLL coated microfluidic chambers grow robustly across the 450µm microgroove, determined by β-III tubulin staining. Treating the neurite growing side (right side from the microgroove) only with increasing concentrations of mixed population of histones isolated from calf thymus results in significantly shorter neurites with the following conditions: Control, Aprotinin-treated, Histones - 5µg/ml, Histones 10(µg/ml) and Histones 20(µg/ml).

**Figure S7.**
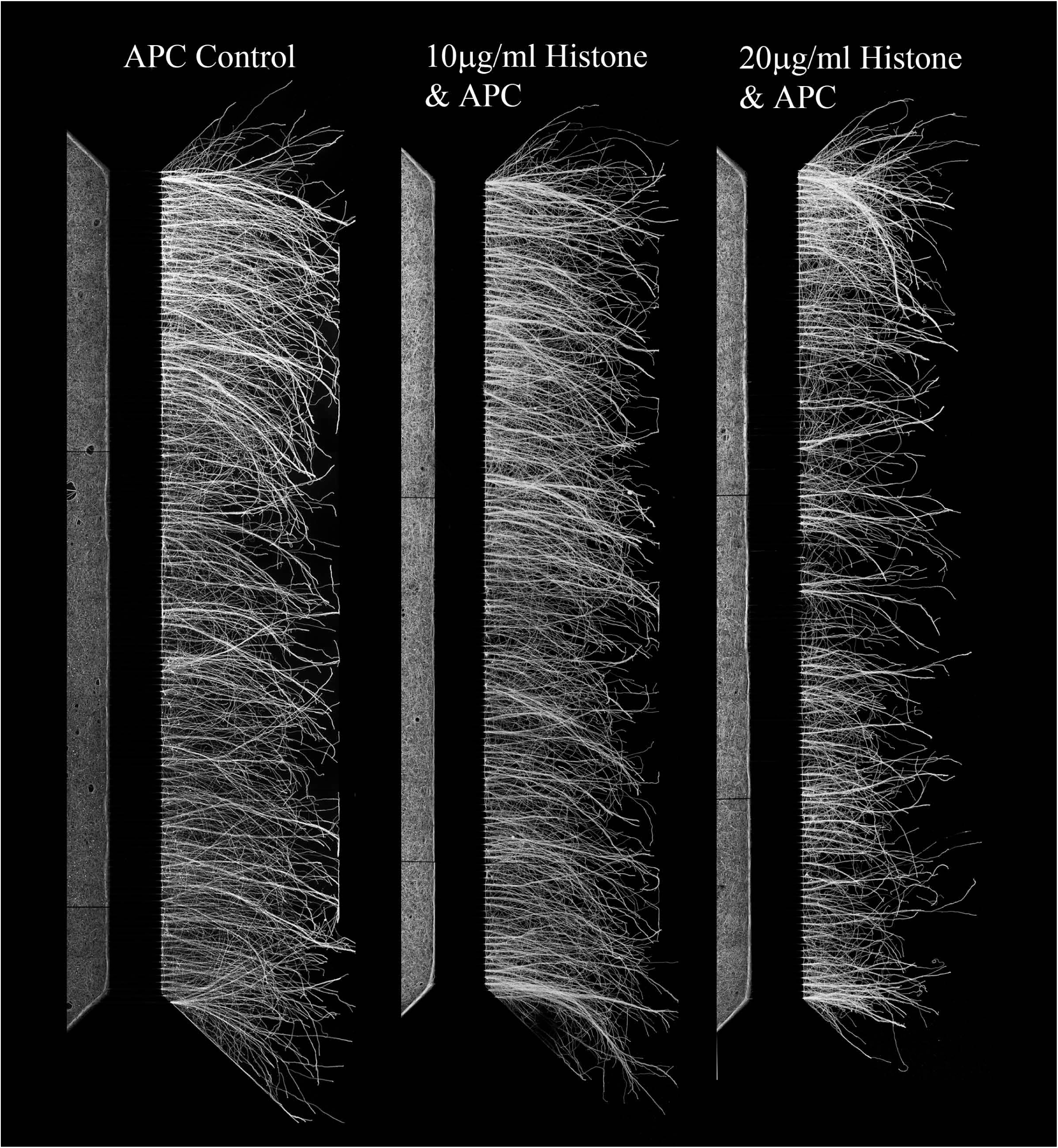
Activated Protein C (APC) blocks the inhibitory effect of Histones. APC alone has no effect on neurite outgrowth, combining APC with histones and applying to the neurite side reverses the inhibitory effect of histones, restoring long neurite outgrowth in microfluidic chambers.

**Figured S8.**
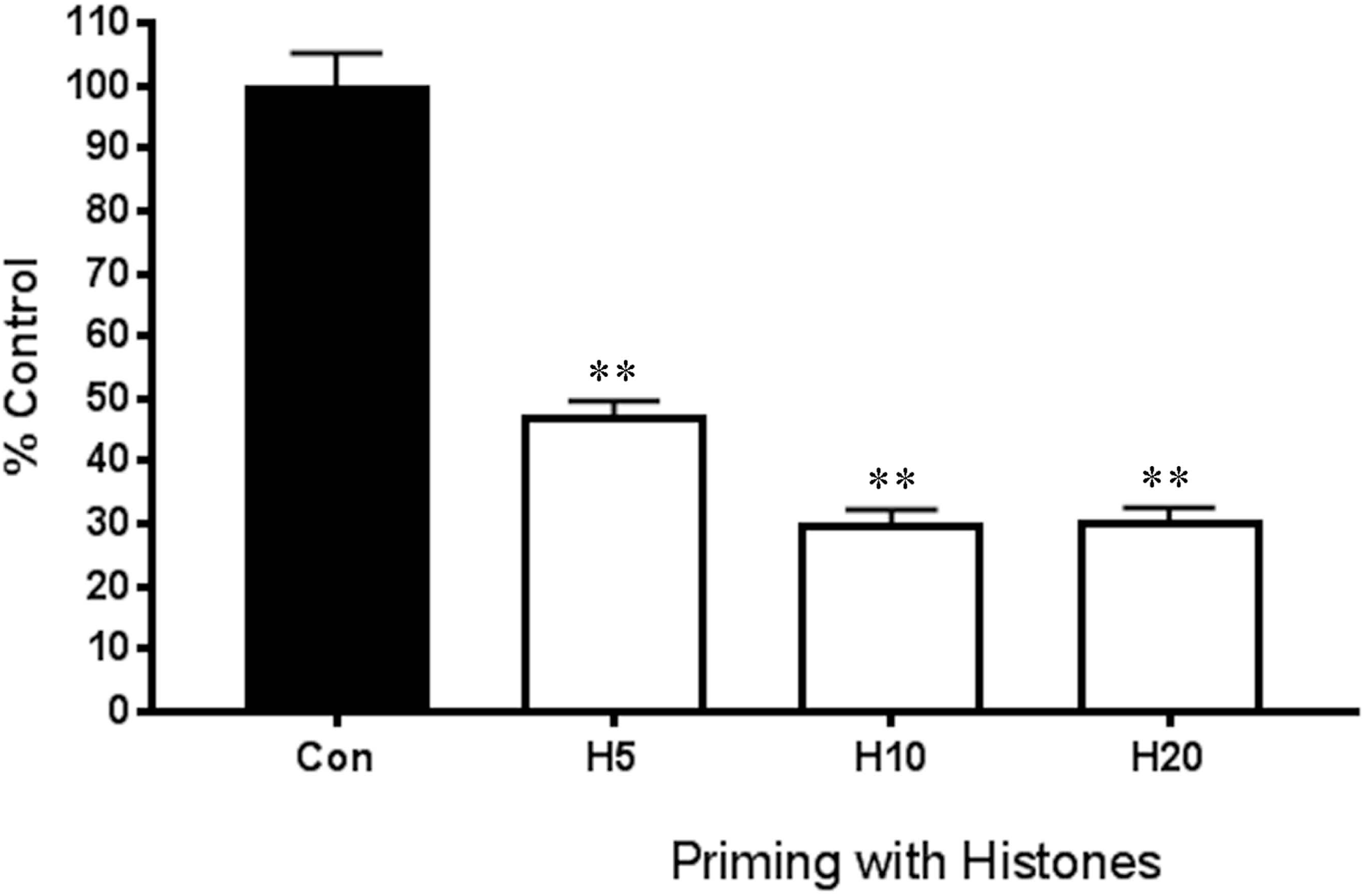
Priming cortical neurons with histones results in significantly shorter neurites. Cortical neurons are treated overnight with either PBS (Con) or 5, 10 or 20µg/ml of mixed prep of histones. The neurons are aspirated and washed with plain NB, trypsinized and plated onto a confluent monolayer of permissive CHO cells in supplemented NB. Note there are no histones subsequently added once on the CHO cells, histones are only present during the overnight priming and then washed out. Once fixed and immunostained with β-III tubulin and quantified, we observed significantly shorter neurites compared to the control (**p<0.01).

**Figure S9.**
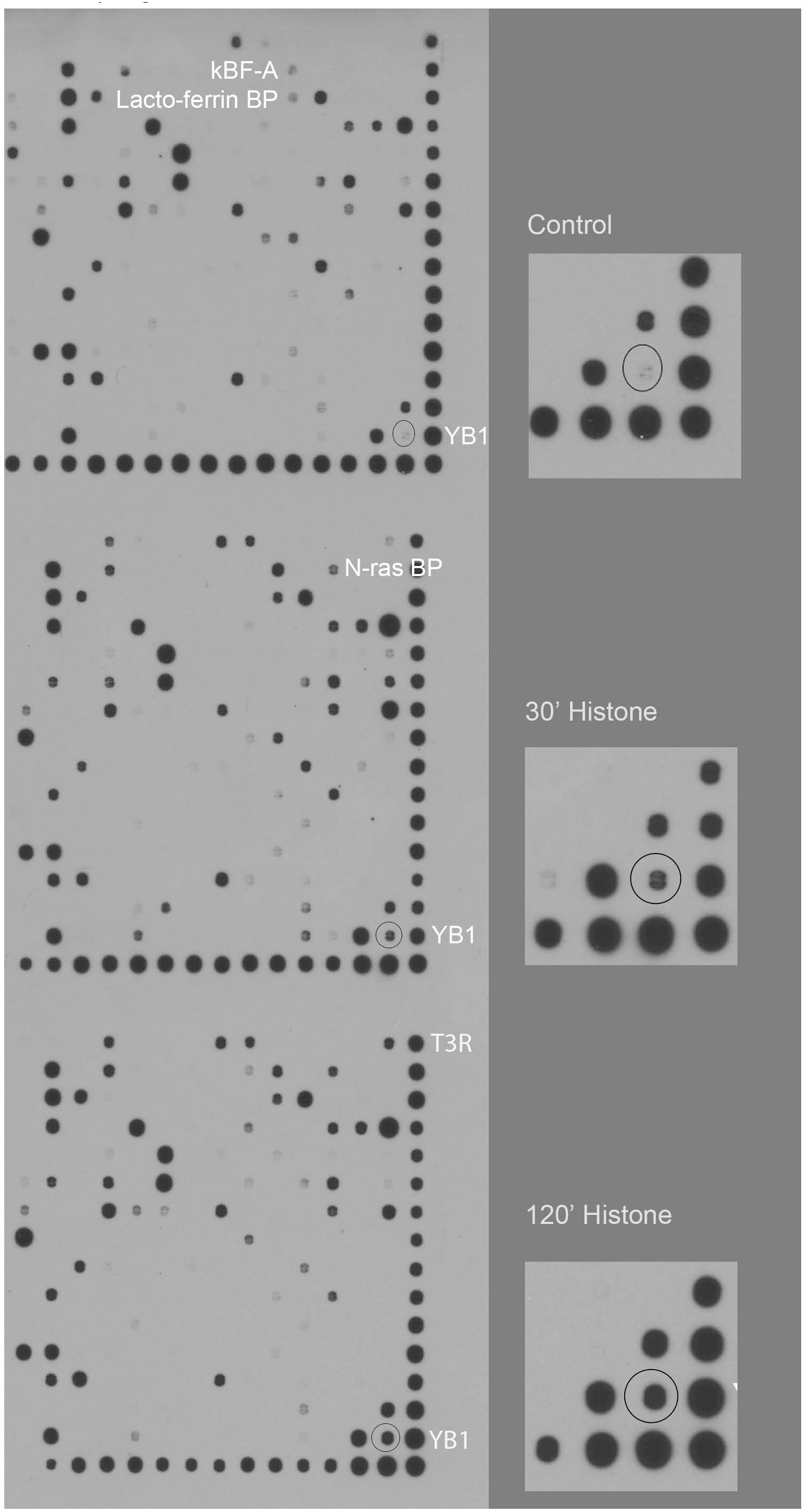
The transcription factor YB-1 is elevated in response to histone treatment on primary cortical neurons. Using a Panomics DNA-protien binding array we observed elevation in the transcription factor YB-1 in primary cortical neurons. We treated our samples with PBS or histones for 30 or 120 minutes before preparing nuclear extracts used in a commercial assay for transcription factor activation. With histone treatment we saw elevation in YB-1 levels as shown in the blowups and circled.

**Figure S10.**
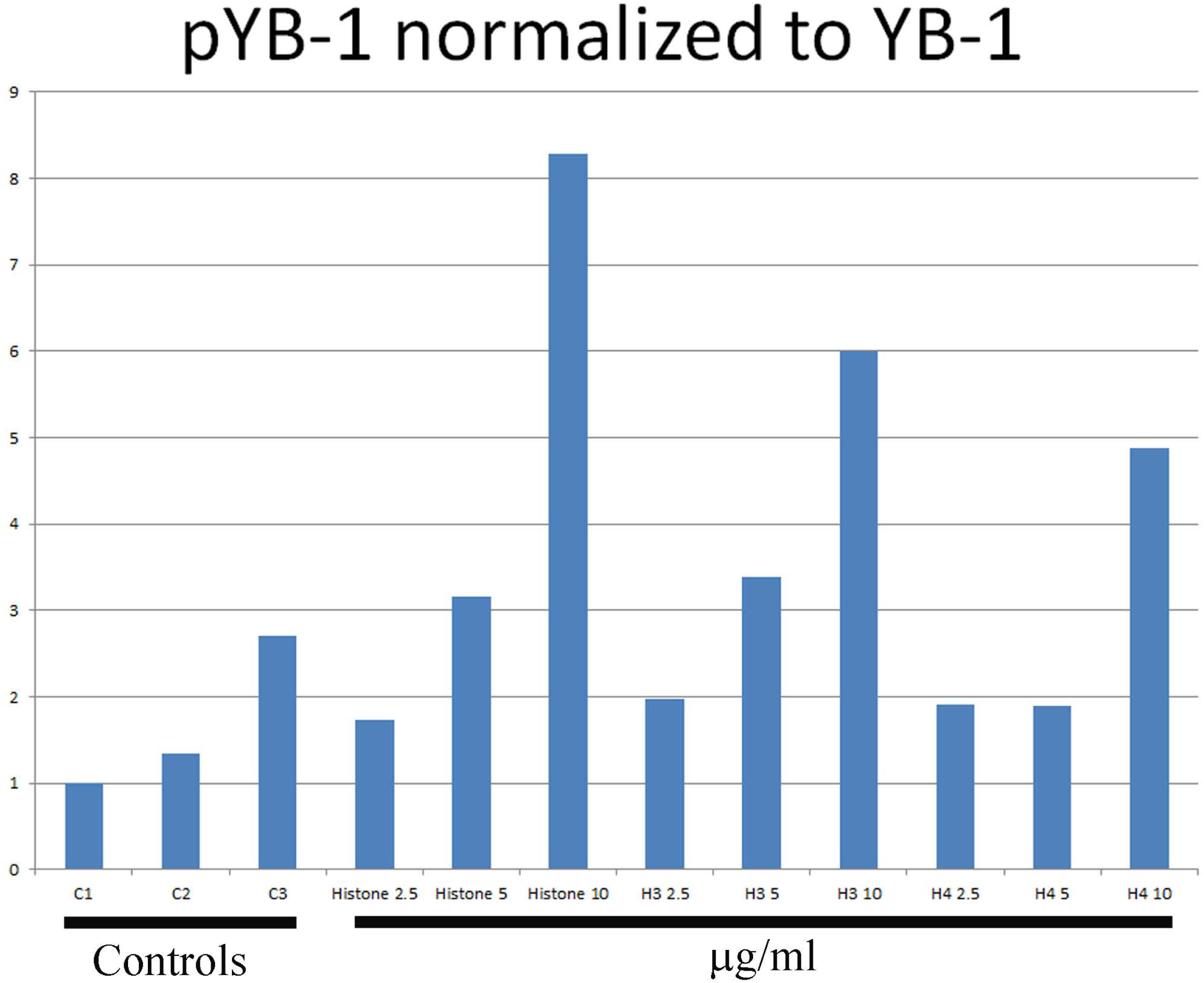
Extracellular histones elevate pYB-1 in cortical neurons. Normalized values from Figure 4B western blots, where we treated cortical neurons for 2hrs with either mixed population of histones (Histone), recombinant H3 or H4 at 2.5, 5 or 10µg/ml. The graph shows normalized values of pYB-1 to total YB-1.

**Figure S11.**
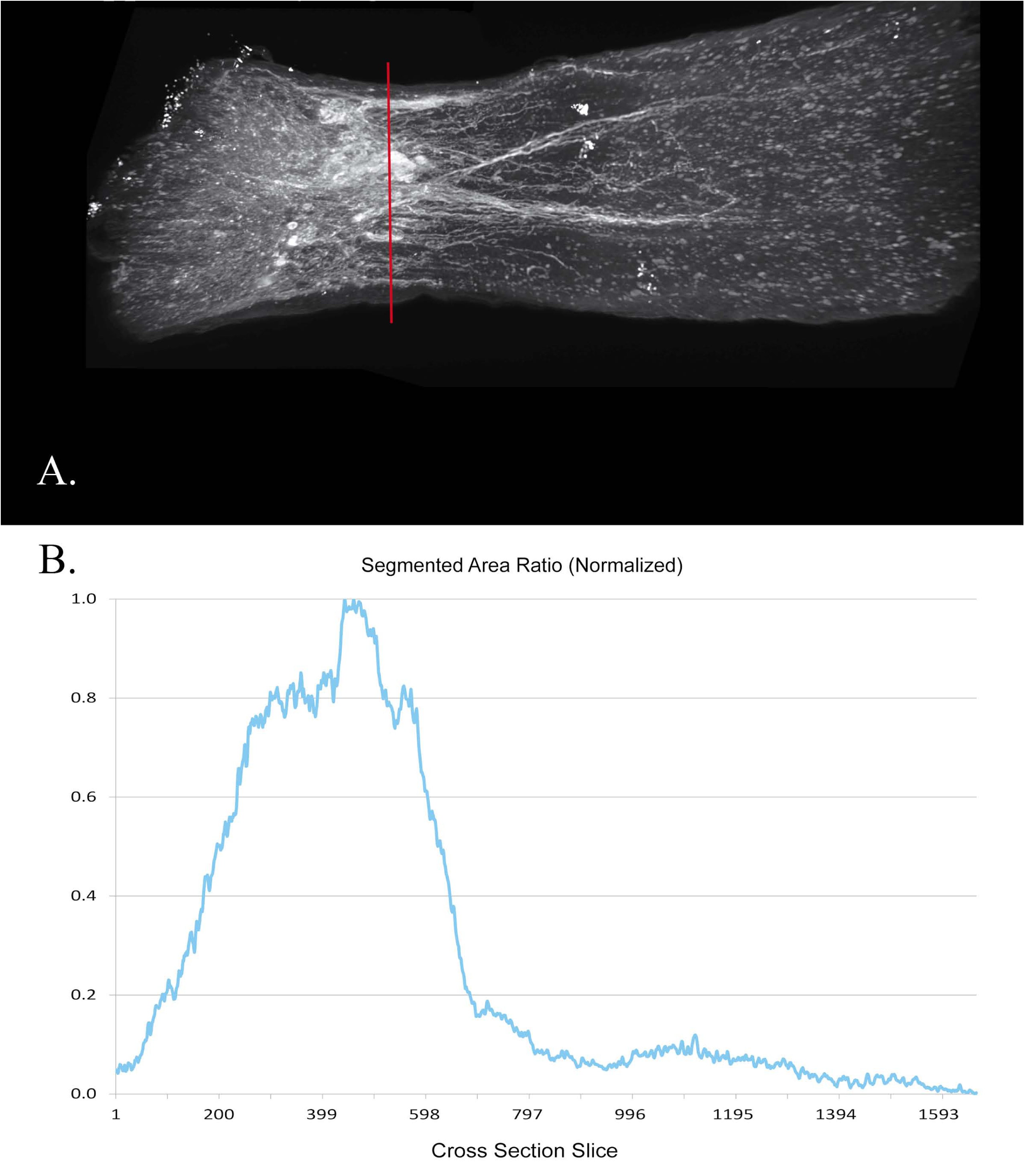
Volumetric analysis in nerves treated with APC after ONC. A. APC-treated nerve, for representation we show how we measure the Normalized Segmented Area Ratio. The cross section slice is set at the middle of the crush site (vertical red bar demarcating). B. The graph representing the Normalized Segmented Area Ratio for the image in A.

**Figure S12.**
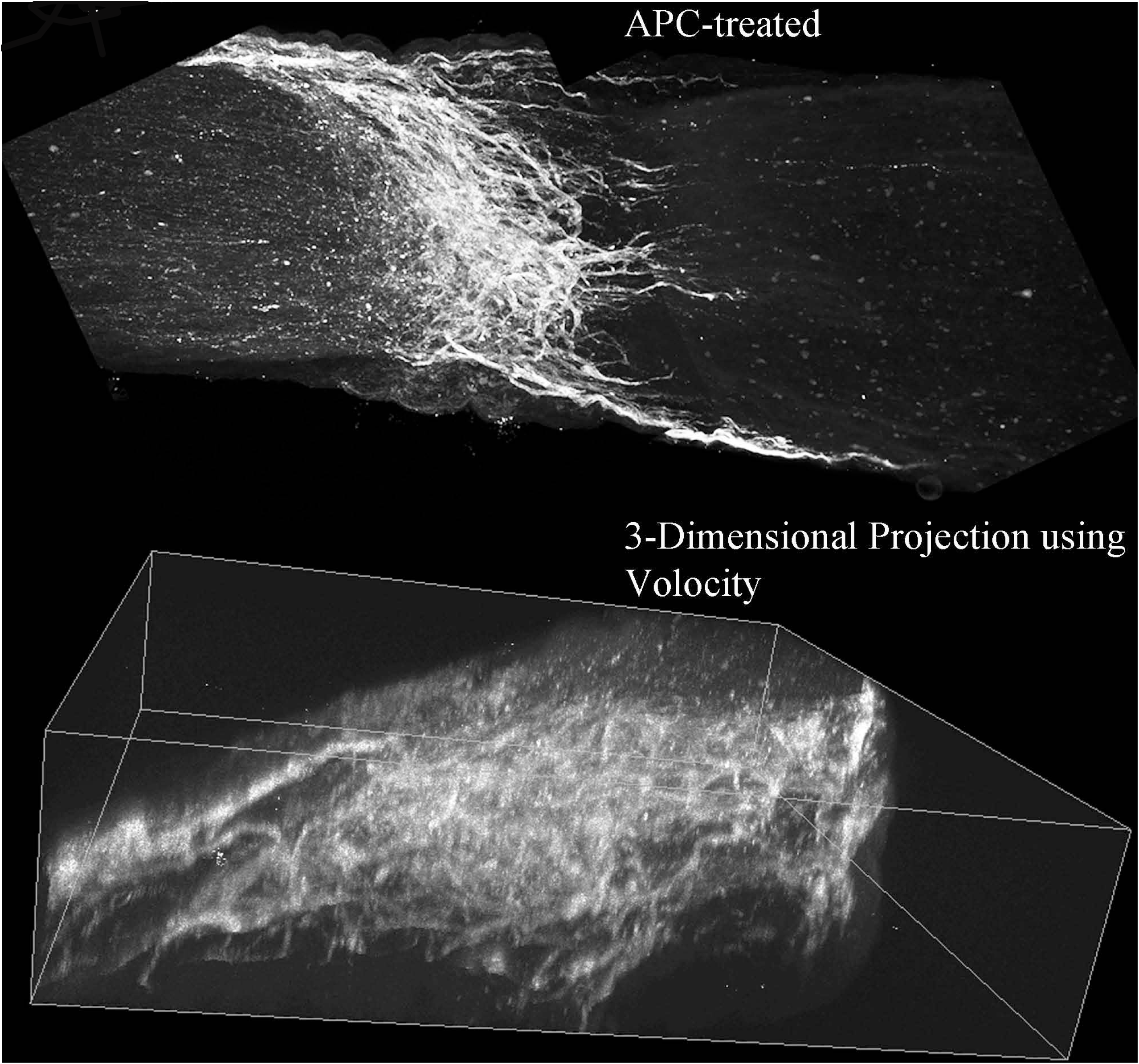
APC treatment in the optic nerve crush using Volocity 3D Projection. Representative ONC treated with APC at the injury site, promoting axonal regeneration. Using Volocity to get a 3D projection of the same nerve to reveal axons regenerating within the nerve.

**Figure S13.**
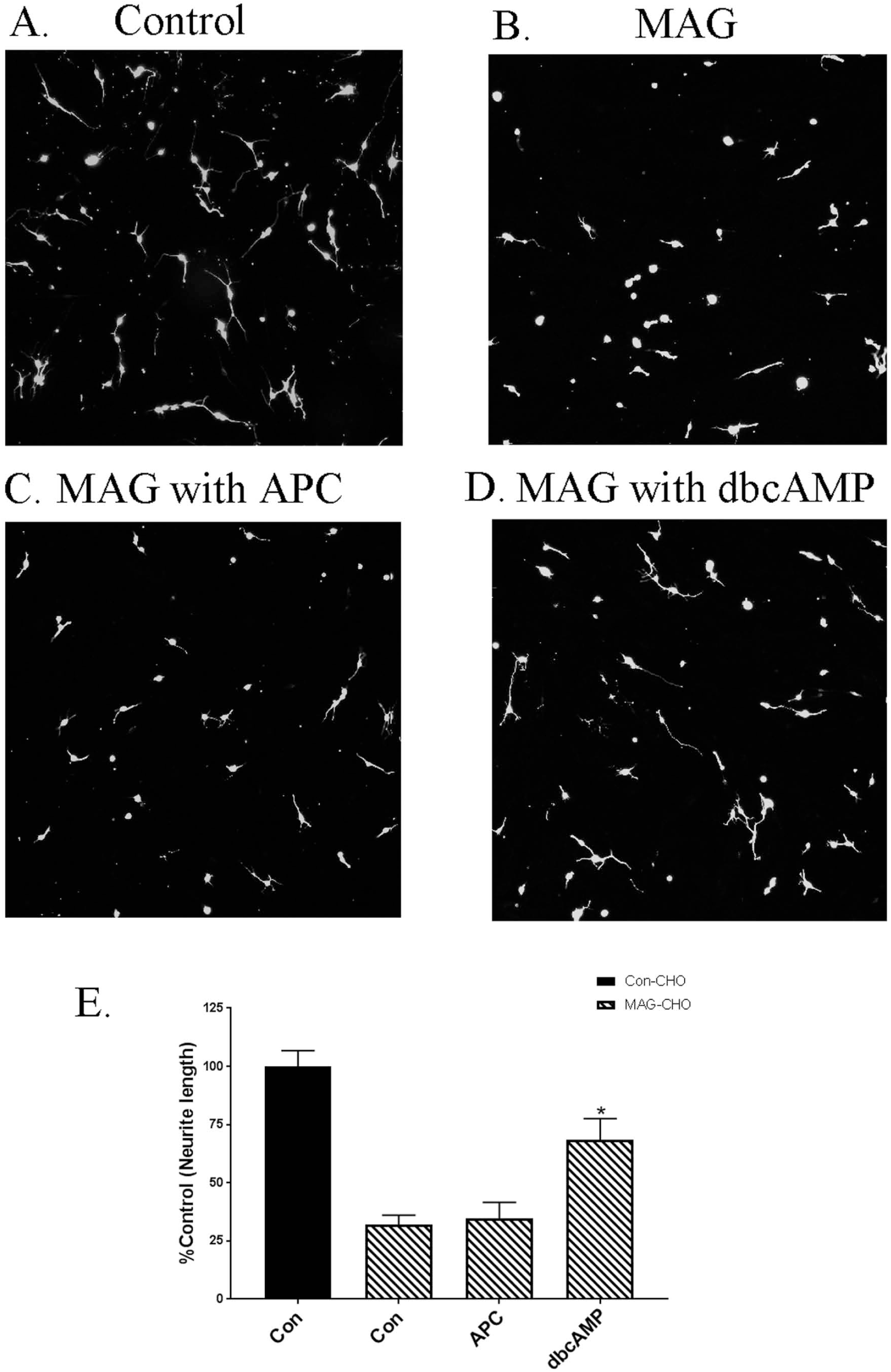
APC does not overcome MAG-mediated inhibition. Cortical neurons are plated on either control CHO cells or Myelin-Associated Glycoprotein (MAG)-expressing CHO cells. On MAG, cortical neurons are inhibited from putting out neurites. APC application to the neurons when plated on MAG does not overcome this inhibition unlike dbcAMP which is known to overcome myelin-mediated inhibitors.

## Methods

### Human CSF fluid from patients with SCI

Human spinal cord injury samples and related ASIA scores from fully deidentified patients were obtained from the International Spinal Cord Injury Biobank (ISCIB), which is housed in Vancouver, BC, Canada. Permission for CSF fluid acquisition was granted by the Clinical Research Ethics Board (CREB) of the University of British Columbia, Vancouver, Canada (Ethics certificate of full board approval H19-00690). Samples of fully deidentified CSF from patients with SCI and normal patients were sent to the laboratory of Ravi Iyengar at the Icahn School of Medicine at Mount Sinai (ISMMS), NYC. The IRB review at ISMMS determined IRB approval was not required for these fully deidentifed CSF samples, as it is not human research as defined by DHHS and FDA regulations.

### Rat primary cortical or hippocampal neuron cultures

Rat primary cortical cultures were dissected from postnatal day 1 Sprague Dawley rat brains. Cortices or hippocampi were incubated twice for 30mins with 0.5 mg/ml papain (Sigma) in plain Neurobasal (NB) media (Invitrogen) with DNase. Papain activity was inhibited by brief incubation in soybean trypsin inhibitor (Sigma). Cell suspensions were layered on an Optiprep density gradient (Sigma) and centrifuged at 1900 x *g* for 15mins at room temperature. The purified neurons were then collected at 1000 x g for 5 mins and counted. Primary cortical neurons were diluted to 35,000 cells/ml in NB supplemented with B27, L-glutamine and antibiotics, and we seeded 300µls of the cells suspension to each well and incubated for 24hours. To quantify the outgrowth we immunostained using a monoclonal antiβIII tubulin antibody (Tuj1;Covance) and Alexa Fluor 568-conjugated anti-mouse IgG (Invitrogen). For quantification, images were taken, and the length of the longest neurite for each neuron was measured using MetaMorph software (Molecular Devices). For western blots primary neurons were lysed in radioimmunoprecipitation assay (RIPA) buffer (50mM Tris-HCl, pH 7.5; 100mM NaCl; 1 % Nonidet P-40; 0.5 % deoxycholic acid; 0.1 % SDS; 1mM EDTA) supplemented with protease and phosphatase inhibitors. After determination of protein concentration, the cell lysates were subjected to immunoblot analysis using standard procedures and visualized by enhanced chemiluminescence (ECL). Polyclonal rabbit antibodies directed against phospho-YB-1 and total YB-1 were from Cell Signaling Technology Inc.

### Rat primary dorsal root ganglion neurons

Rat dorsal root ganglion (DRG) neurons from extracted from P5-P7 day old rat pups. The neurons were first treated with with 0.015% collagenase in Neurobasal-A media for 45 min at 37°C. This was followed by a second incubation in collagenase for 30 min at 37°C, with the addition of 0.1% trypsin and 50 µg/ml DNase I. Trypsin was inactivated with DMEM containing 10% dialyzed fetal bovine serum, and the ganglia were triturated in Sato’s media.

### Rat primary astrocytes

We prepared mixed cortical cultures from P3-P5 pups. The cortices were removed and the meninges carefully peeled away. They were treated with 0.1% trypsin and 50 µg/ml DNase I for ten minutes at 37°C. Trypsin was inactivated with DMEM containing 10% dialyzed fetal bovine serum (FBS). We triturated the tissue and strained the cell suspension which was seeded into a pre-coated PLL T75 tissue flask in the growth media of DMEM with 10%FBS. Media was replaced within 24hrs and then every 3 days until we had a confluent layer of cells (10-14 days). To enrich for astrocytes, the media was replaced with plain NB (no supplements but containing Hepes) and we shaked the culture at 200RPM at 37°C. The media was aspirated off and we added fresh DMEM with 10%FBS and returned to the incubator overnight. The next day we would trypsinize the remaining cells for for ten minutes at 37°C, or longer depending on how the cells are lifting off the flask. Trypsin was inactivated with DMEM containing 10% dialyzed fetal bovine serum, and the astrocytes were counted.

### Neurite outgrowth assay

Monolayers of control or MAG-expressing Chinese hamster ovary (CHO) cells or primary astrocytes were plated on eight well chamber slides as previously described (30). Purified P1 hippocampal, P1 cortical, P5–P6 CGN, or P5–P6 DRG rat neurons were diluted to 35,000 cells/ml in Sato’s media and treated with either 1 mM dbcAMP (Calbiochem), mixed population or recombinant Histones (1-20µg/ml) or with APC (1-20µg/ml). Neurons were incubated for 14–18 h at 37°C and immunostained using a monoclonal anti-βIII tubulin antibody (Tuj1;Covance) and Alexa Fluor 568-conjugated anti-mouse IgG (Invitrogen). For quantification, images were taken, and the length of the longest neurite for each neuron was measured using MetaMorph software (Molecular Devices). *Microfluidic neurite outgrowth assay-*Square microfluidic chambers (150 or 450 microgroove) were purchased from Xona microfluidics. The chambers were sterilized under UV for 15mins and soaked in 70% ethanol for 2mins and allowed to air dry under a sterile TC hood. We used MatTek dishes (P50G-1.5, MatTek corp.) that we pre-coated with PLL overnight, rinsed 3 times with sterile water and air dried overnight in a sterile hood. Using sterile forceps we carefully place the microfluidic chamber on the glass area of the MatTek dish, and gently apply pressure to ensure the chambers are sitting on the glass. 3 whole brain P1-2 cortices after papain enzymatic digestion and Optiprep gradient were resuspended in 200µls of NB with full supplements. We then seeded 15µls of the neuronal suspension on one side of the microgroove and place in a 37°C incubator for 20mins to allow the neurons to adhere. All wells were filled with 150µls of supplemented NB. Once we observe neurites growing across the microgroove (2-3days), we then did treatments on the neurite compartment and wait 48hrs before we add 4% Paraformaldehyde with 4%Sucrose. We used the Neon transfection (Invitrogen) method to overexpress YB-1.

### Rho Activation Assay

Rat cortical neurons plated in 10cm^2^ dishes (approximately 10 million neurons per dish) were used with a commercially available Rho Activiation Assay Kit (Millipore). The neurons were placed in plain NB media for 4hrs prior to assay. In brief, following the manufacturers protocol, we lysed the cortical neurons on ice with MLB buffer supplemented with anti-protease and anti-phosphatase cocktails (Calbiochem) and spun the lysates at 14,000 x g for 5 mins. The supernatant was collected and added to 35µls of agarose beads coupled to Rhotekin Rho Binding Domain and rocked in the cold room for 45mins, a small sample of each supernatant was not added to the beads and was used as total Rho loading controls. Beads were washed three times with MLB buffer and 20µls 2xLaemmeli buffer was added and samples were prepared for western blotting. Proteins were separated on pre-cast 420% gradient gels (Thermo Fisher Scientific) and transferred to nitrocellulose at 75V for 1 hr. Membranes were successfully probed with rabbit anti-RhoA (1:1000; Cell Signaling Technology) and HRP conjugated anti-rabbit IgG (1:2000; Cell Signaling Technology). Membranes were reacted with Pierce ECL Western Blotting Substrate or SuperSignal West Femto Maximum Sensitivity Substrate (Thermo Fisher Scientific). Densitometirc measurements were made using NIH Image J software.

### Optic nerve crush experiments

Adult male or female Sprague-Dawley or rats (250–280 g, approximately 8-10 weeks old) were anesthetized with isoflurane and placed in a stereotaxic frame. The right optic nerve was exposed and crushed with fine forceps for 10s. We placed gelfoam either soaked in PBS or with APC [4.1mg/ml] (Haematologic Technologies, Inc) were placed over the injury site. 3 days prior to sacrificing we label the regenerating axons with 5µls of 1mg/ml Cholera Toxin B (CTB) coupled to Alexa-488 which we intravitreally inject. Animals were anesthetized with ketamine (100mg/kg) and xylazine (20mg/kg) injected intraperitoneally and transcardial perfusion with 4% PFA after a 14 days postsurgical survival period. When animals were deeply anesthetized we transcardially perfused with cold 4%Paraformaldehyde (PFA) in PBS, pH 7.4. The optic nerves and chiasm attached were dissected out and post-fixed in 4%PFA overnight at 4°C, rinsed for one hour in PBS and then we prepare them for chemical clearing. Since the advent of the 3DISCO clearing techniques, we are no longer dependent on sectioning the tissue, this method also eliminates bias associated with artifacts produced by sectioning. The whole nerve is placed in a graded series dehydration technique in Tetrahydrofuran (THF;Sigma) diluted in water: 50%,70%,80%,100% and 100% again. Each THF dilution is for 20mins at room temperature on an orbital shaker (15). Followed by Dicholoromethane (Sigma) for 5mins at room temperature on an orbital shaker and followed by the clearing agent, Dibenzyl ether (DBE; Sigma) overnight at room temperature on an orbital shaker. Microscope slides are mounted with Fastwell chambers (Electron Microscopy Sciences) and the cleared sample is place on the slide and covered with DBE, a No. 1.5 micro cover glass is used the cover the sample. We image the whole sample on an Olympus Multiphoton microscope with a 25X water immersion lens. All procedures were approved by the IACUC of the Icahn School of Medicine at Mount Sinai in accordance with NIH guidelines. Multiphoton microscopy was performed in the Microscopy CORE at the Icahn School of Medicine at Mount Sinai and was supported with funding from NIH Shared Instrumentation Grant 1S10RR026639-01.

### Multiphoton Image Analysis

Images were deconvolved using AutoQuant X Software (Media Cybernetics). Deconvolved images were then analyzed using Amira Software version 6.0.1 (Thermo Scientific). Z-stacks were uploaded to create a 3D volume rendering of the crush site and labelled neurites. For samples containing more than one Z-stack, image stacks were aligned and merged using Amira’s merge module. The images were then rotated to position the crush site proximal to the retina on the left with regenerated neurites growing away from the crush site towards the right. The transformed data was resampled and axes swapped such that the XY plane became the cross section along the length of the nerve. Images were cropped if needed to exclude any partial cross sections of the ends of the nerve caused by image rotation. Labelled regenerated neurites were segmented by thresholding pixels using Amira’s Segmentation Editor. The volume ratio of segmented neurites over total nerve volume was measured for each sample. Study results were then normalized and plotted as a bar graph. The ratio of segmented area over total nerve area for each cross section was normalized and plotted as area ratio per slice. Area ratio plots were created starting from the behind the crush site proximal to the retina and following along the length of the nerve for the segmented regenerated neurites.

### Cell fractionation and Panomics transcription factor activation arrays

Nuclear and cytosolic extracts were isolated from cortical neurons following instructions from a commercial kit (Pierce) (19,31). The supernatant containing the nuclear extract was used in a Panomics Combo Protein-DNA Array (Affymetrix, MA1215) to study transcription factor activation following the manufacturer’s protocol. The array was visualized by enhanced chemiluminescence and determined where we had a positive signal.

### siRNA transfection

All siRNA were purchased from Accell smartpool siRNA (Thermo Scientific), following the manufacturers protocol. The following siRNA for rat were utilized: Rtn4r (NgR, EntrezGene 113912), toll-like receptor 2 (EntrezGene 310553) and toll-like receptor 4 (EntrezGene 29260), scrambled non-targeting siRNA was also utilized for controls. Cortical neurons were plated on one side of the microfluidic chambers. We waited 2-4 hrs after plating to allow the cells to adhere. Utilizing 1µM siRNA in 150µls of Accell Delivery Media was added to the neuronal cell bodies and incubated overnight at 37C incubator. The next morning, we added 150µls of supplemented NB. 48hrs after siRNA-delivery, we added our histones to the neurite side of the chambers. We waited 48hrs before stopping the experiment with 4% paraformaldehyde and 4% sucrose. We then stained the chambers with β-III tubulin and quantified the neurite length.

### Statistical analyses

All analyses were performed using GraphPad Prism software, and data are represented as mean ± SEM. Statistical significance was assessed using paired one-tailed Student’s *t* tests to compare two groups, and one-way ANOVAs with Bonferroni’s *post hoc* tests to compare between three or more groups.

### Bulk Sequencing Molecular Preparation

We used primary rat cortical neurons for mRNA sequencing, total RNA was extracted using Trizol (Thermo Fisher). All samples RNA integrity was checked by Agilent 2100 Bioanalyzer and all samples had RIN value ≥9 (31). mRNA libraries for RNA sequencing were constructed using the Truseq stranded mRNA kit (Illumina Inc.) which converts the mRNA in a total RNA sample into a library of template molecules of known strand origin. It captures both coding and non-coding RNA which are poly-adenylated. Briefly, poly-T oligos bound to magnetic beads are used to pull down the poly-A containing mRNA molecules from total RNA and unbound contaminants are washed away. The mRNA is then eluted from the beads and fragmented into smaller pieces using divalent cations under elevated temperature to desired insert sizes depending on the original RNA integrity and required sequencing reads configuration. The resulting mRNA fragment inserts are then reverse transcribed to first strand cDNA using reverse transcriptase and random primers. Use of Actinomycin D during first strand synthesis prevents spurious DNA-dependent synthesis and drives a RNA-dependent synthesis. Strand specificity is achieved during second strand cDNA synthesis by replacing dTTP with dUTP in the Second Strand marking mix which includes the DNA Polymerase I and RNase H. Incorporation of dUTP in second strand synthesis quenches the second strand during amplification. 3’ ends are then adenylated to prevent them from ligating to each other during the adapter ligation reaction. Unique dual index adapters (i5 and i7) are then ligated allowing for greater sample indexing diversity and prepares the ds cDNA for hybridization onto a flow cell. The indexed double stranded cDNA was then enriched with PCR and purified to create the final cDNA library ready for quantification prior to loading on the flow cell for sequencing.

### Data Pre-processing Prior to Bulk Sequencing Differential Expression Analysis Pipeline

#### RNASeq and downstream analysis

To reduce experimental artifacts caused by read imbalances we downsampled the sequencing reads of each sample to the number of reads that were detected in the sample with the lowest read counts, as described previously (32). Downsampled reads were aligned to the rat reference genome ’rn6’ using the ensemble annotation and STAR 2.5.4b (33). Differentially expressed genes were identified with cufflinks 1.3.0 (34) (FDR 5%, minimum log_2_(fold change) = +/- log_2_(1.3)). Up- and downregulated genes were subjected to pathway enrichment analysis using Wikipathways and Gene Ontology biological processes, downloaded from the Enrichr website (35,36), as described previously (37).

