## Supplementary material for "Extracellular histones, a new class of inhibitory molecules of CNS axonal regeneration": Full Supplement

### Supplementary Figure 1A

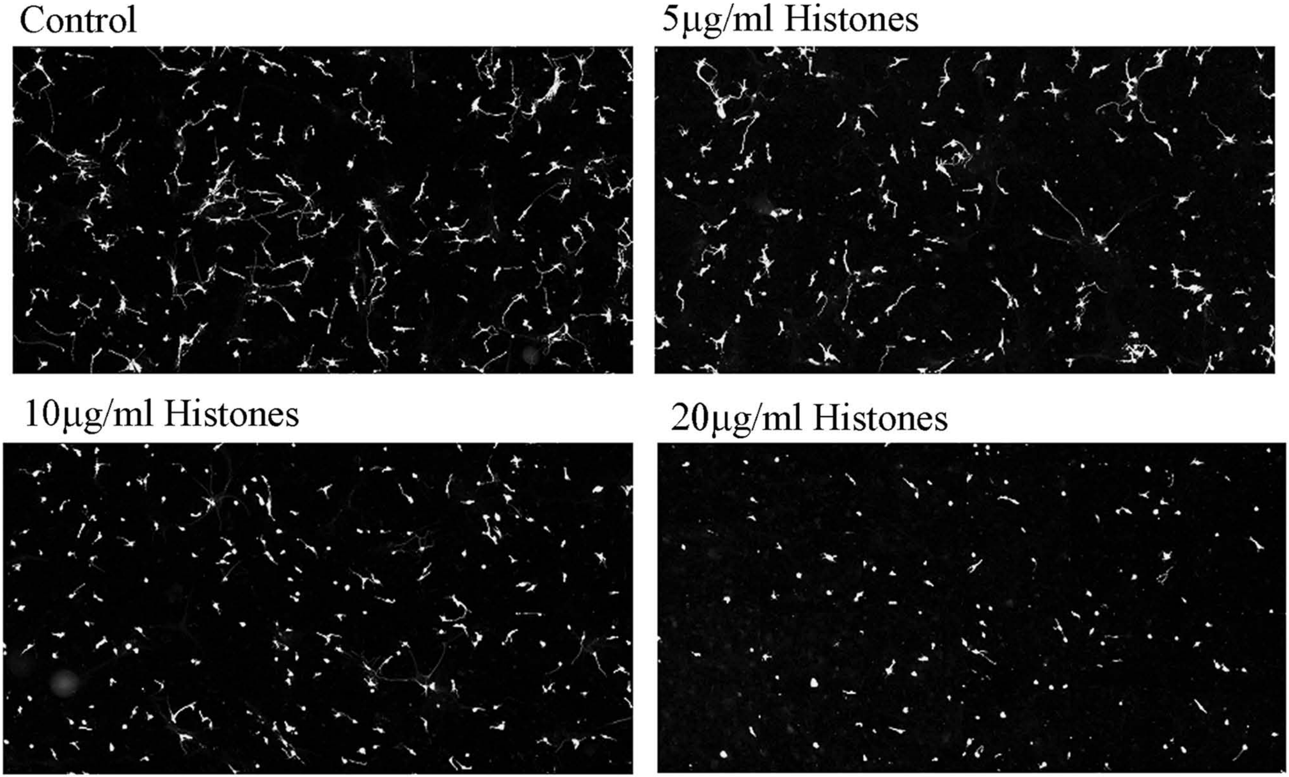

Supplementary Fig 1B

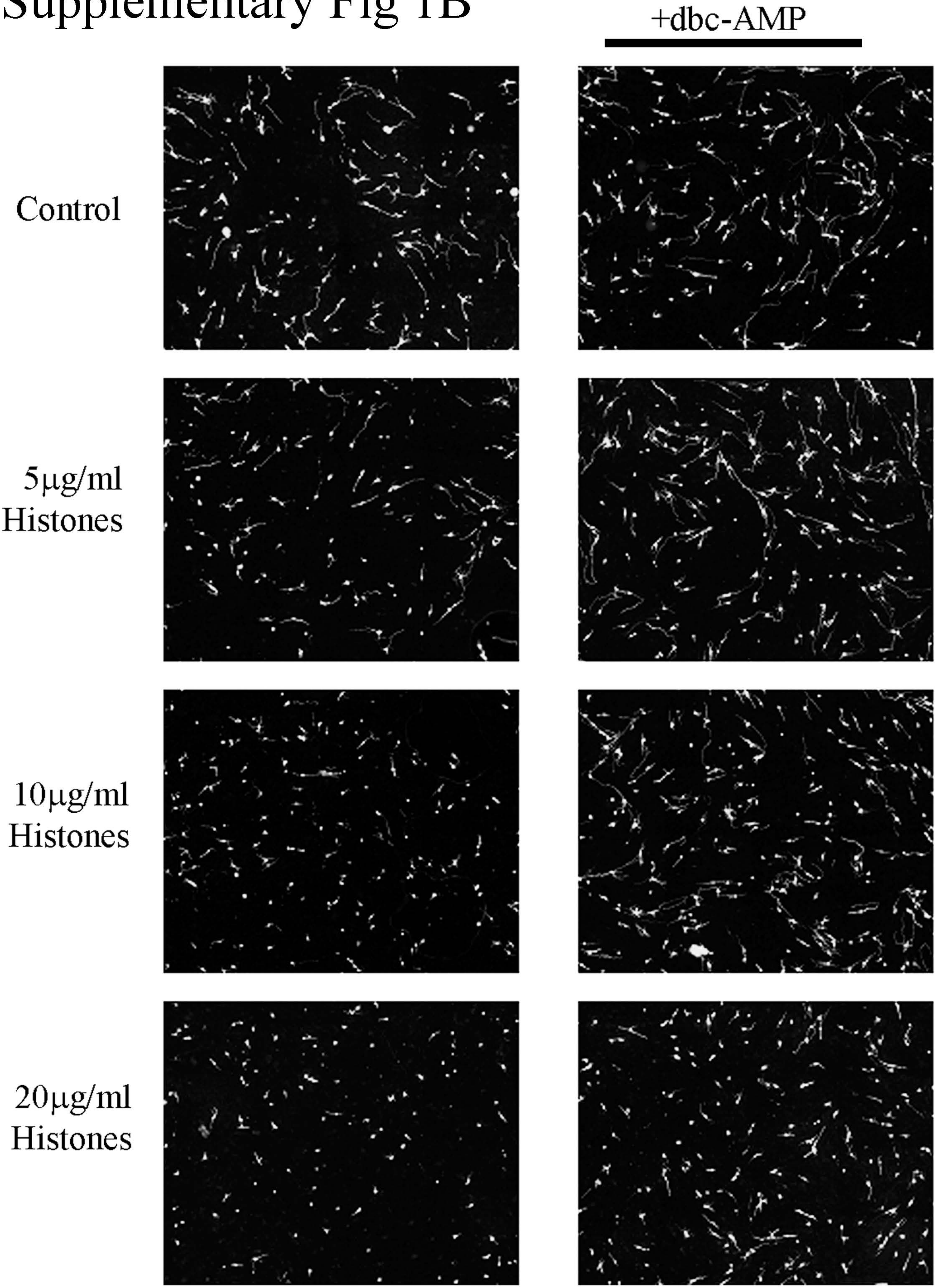

### Figure S1C

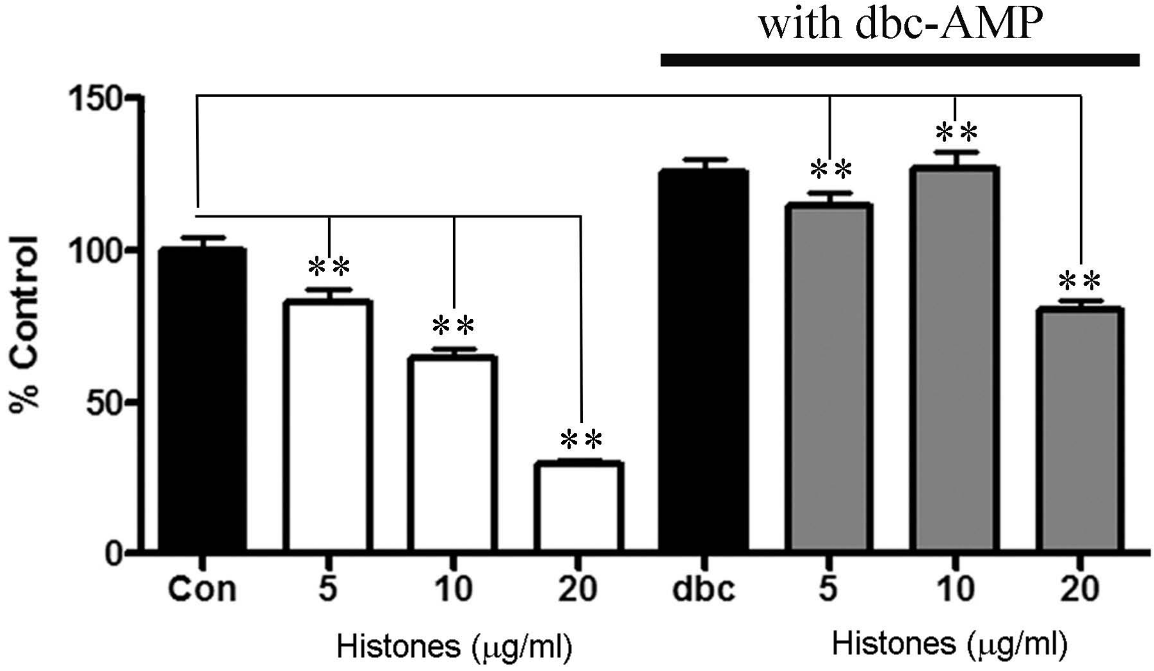

with dbc-AMP
150
100
50
0
% Control
**
**
**
**
**
**
Con
5
10
20
dbc
5
10
20
Histones (μg/ml)
Histones (μg/ml)

### Supplementary Figure 2

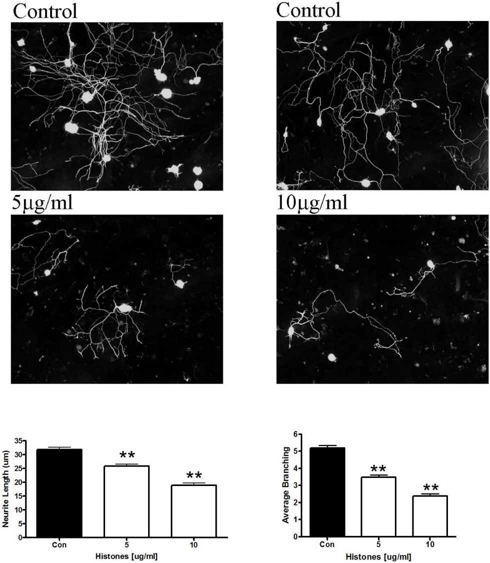

S3A.

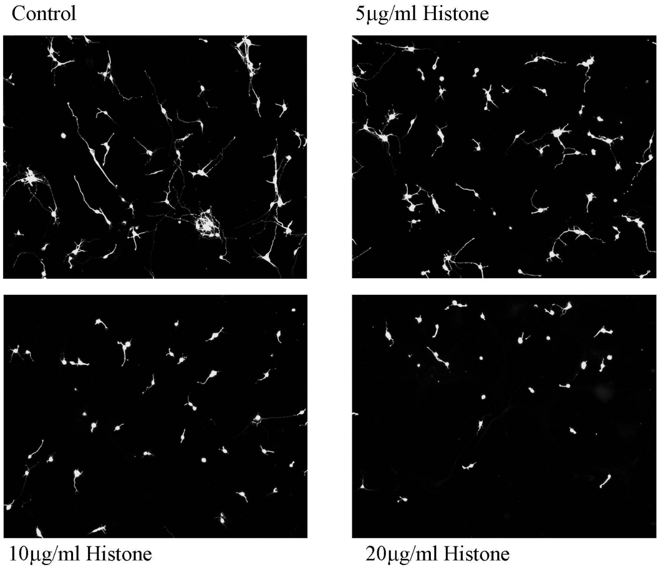

Control
5μg/ml Histone
10μg/ml Histone
20μg/ml Histone


S3B.

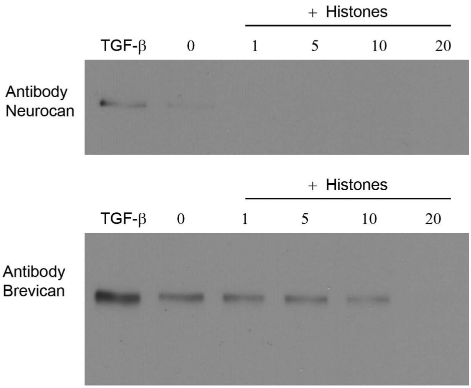

+ Histones
TGF-β
0
1
5
10
20
Antibody Neurocan
+ Histones
TGF-β
0
1
5
10
20
Antibody Brevican

Supplementary Figure 4

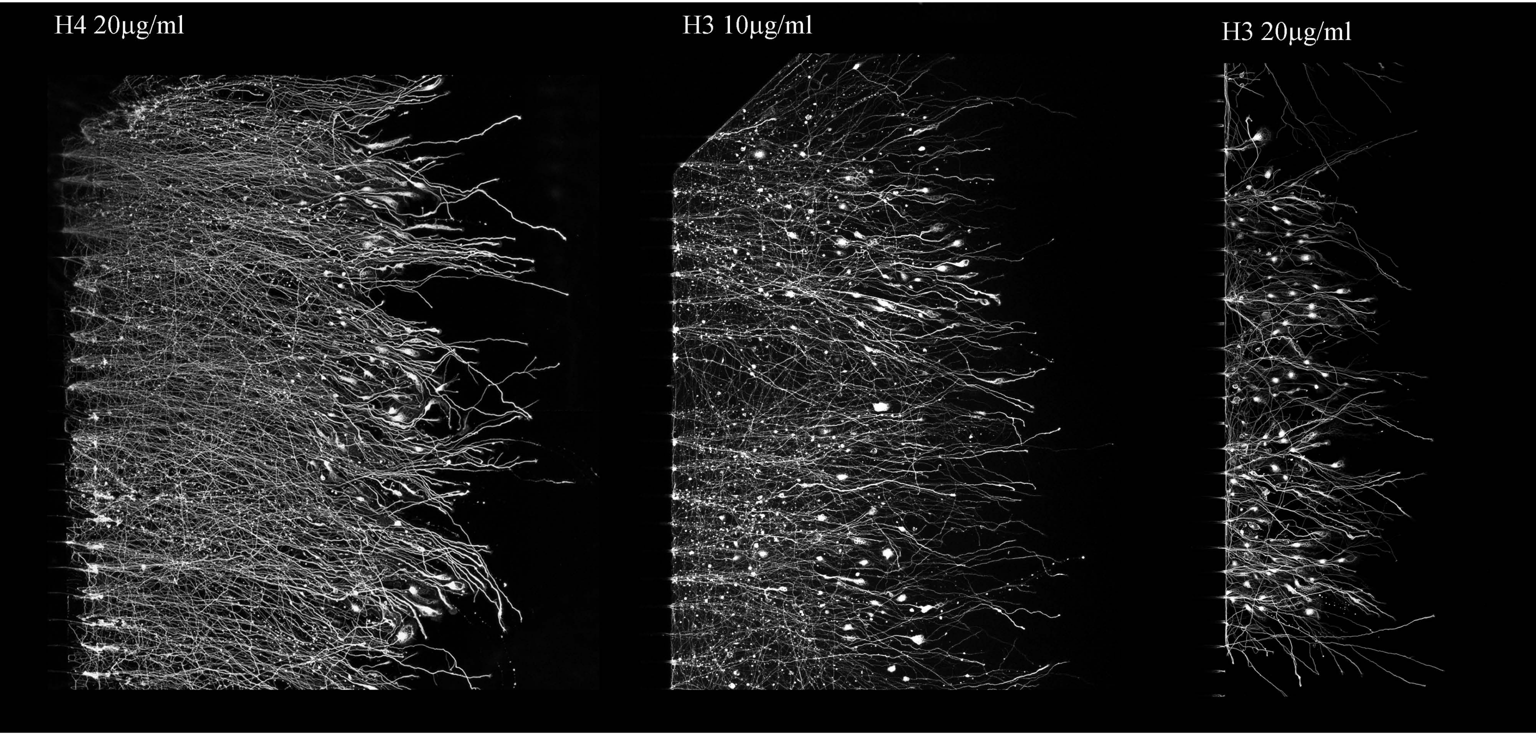

### Supplementary Figure 5

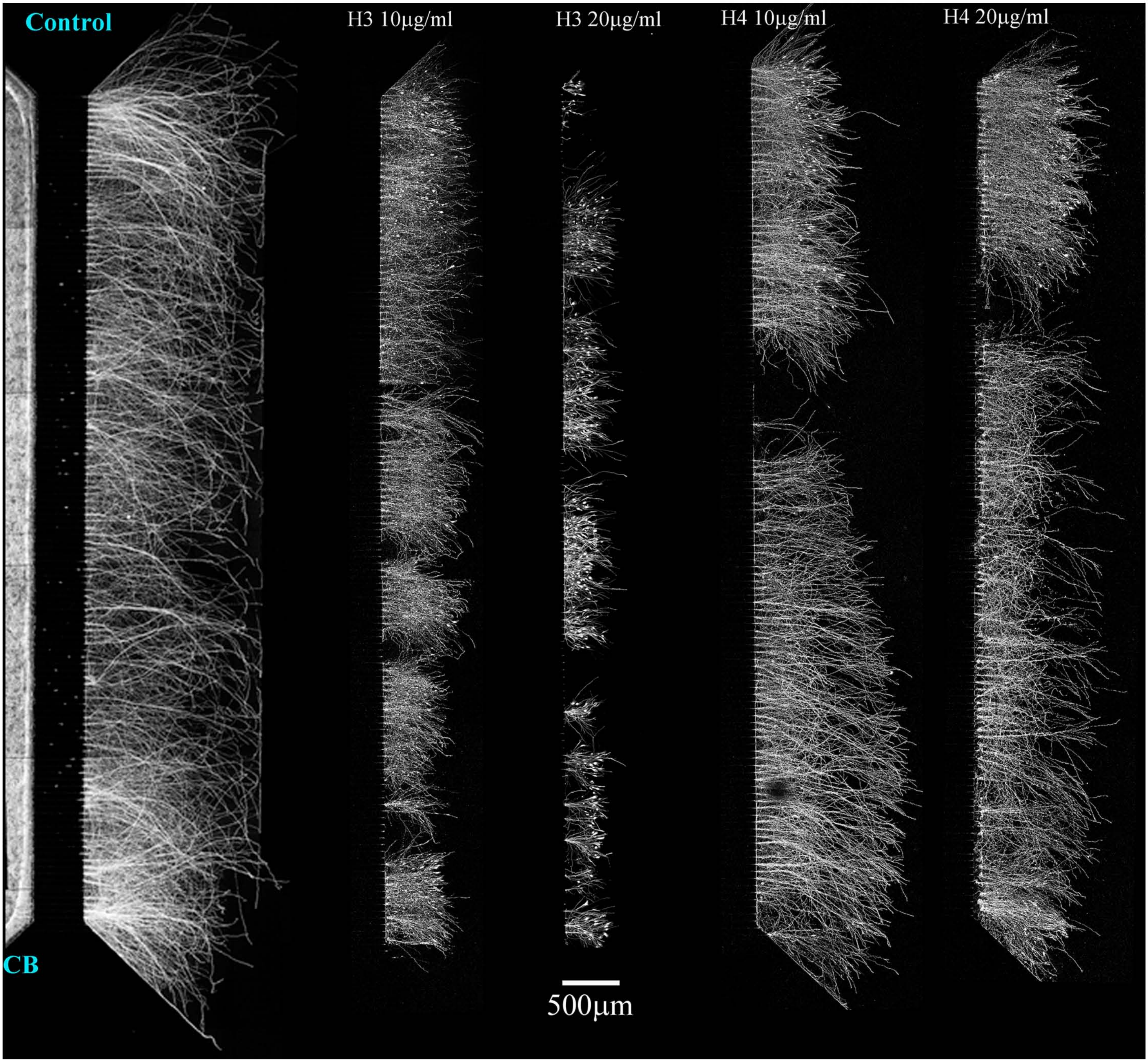

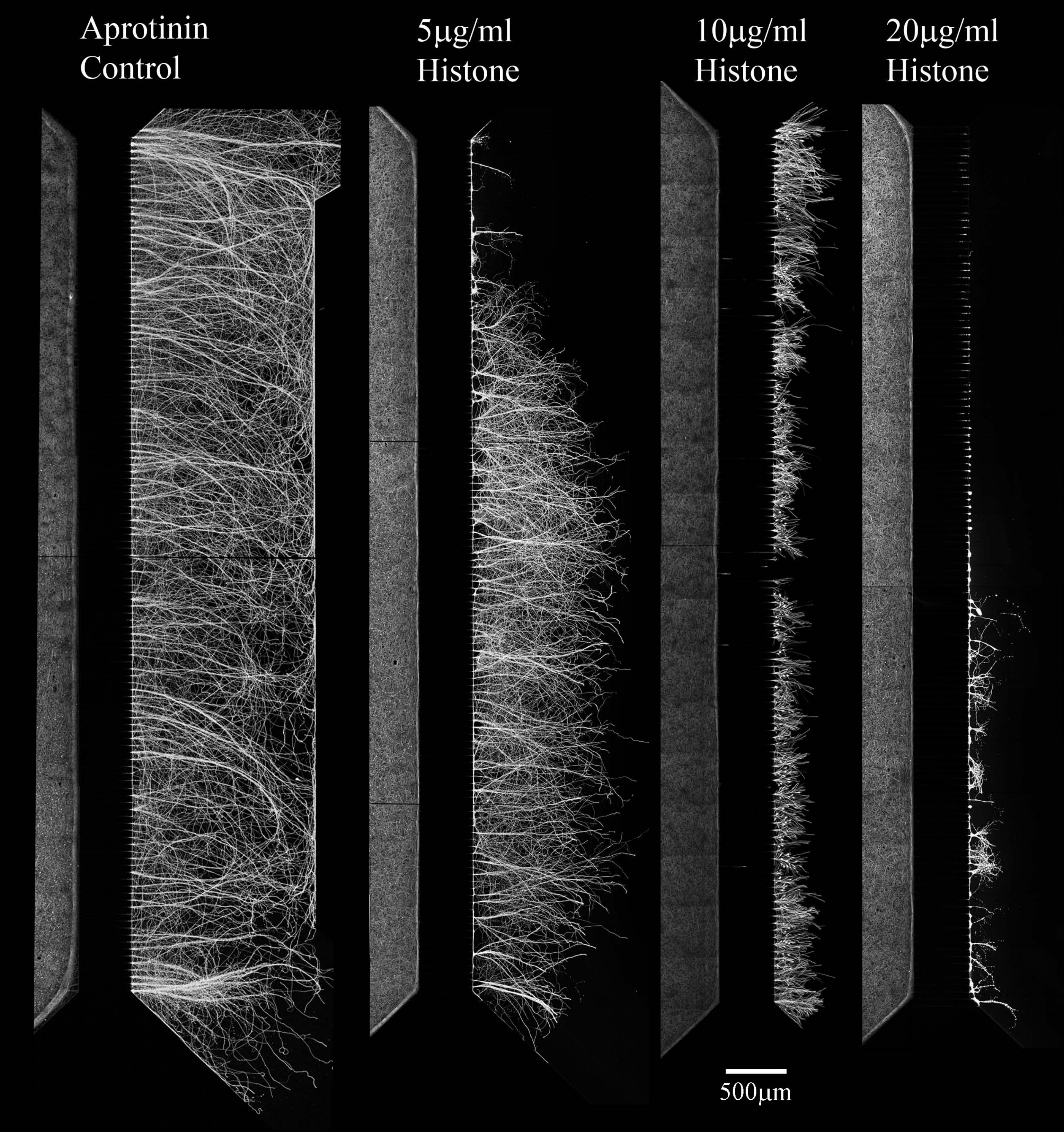


Supplementary
Figure 6

### Supplementary Figure 7

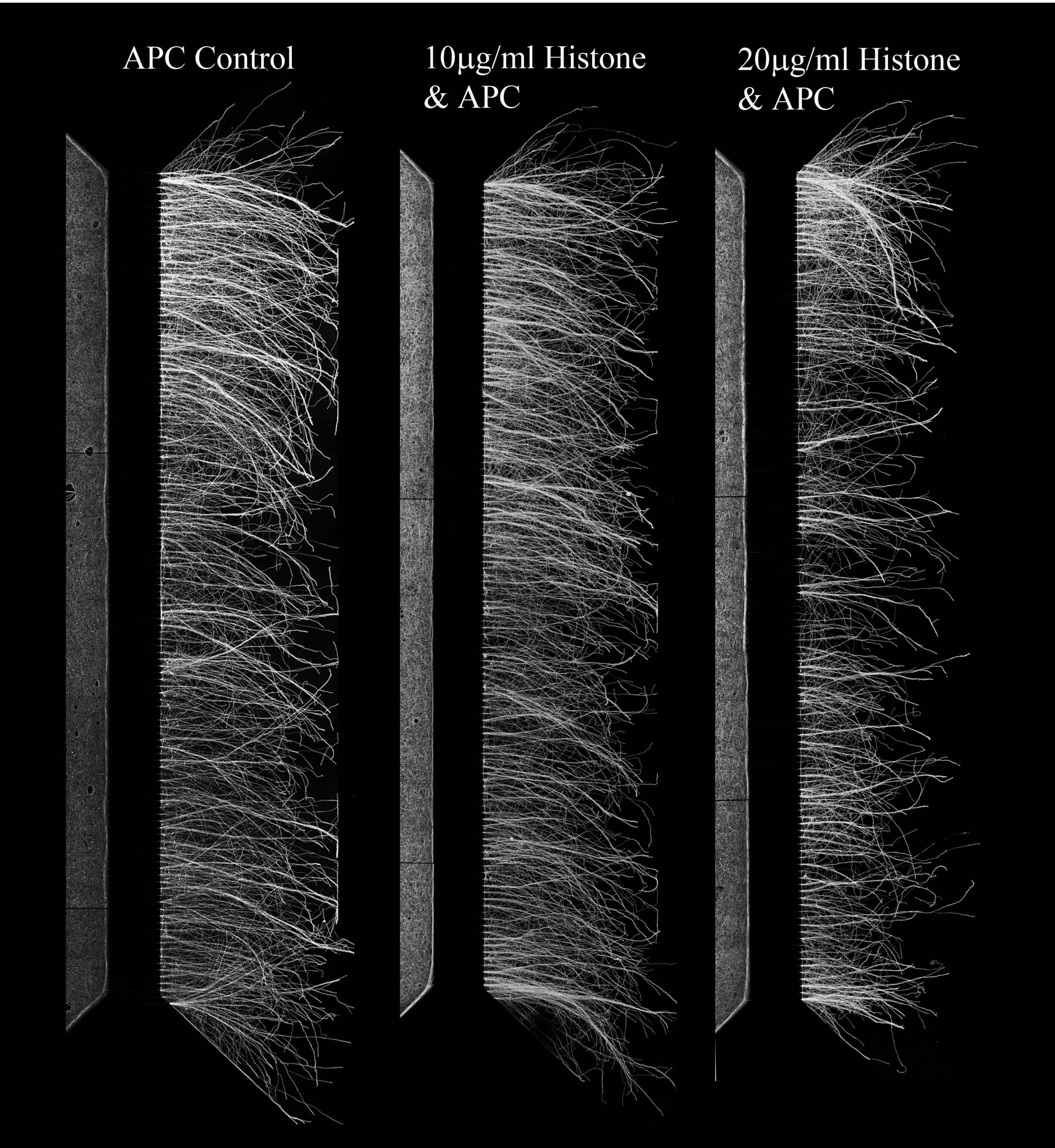

Supplementary Figure 8

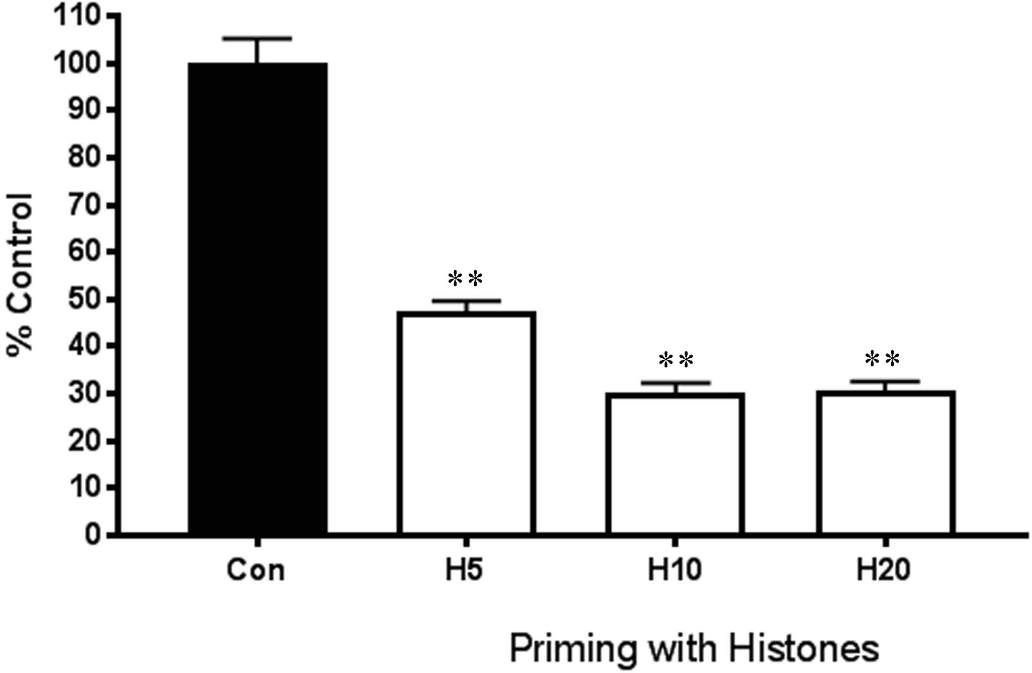

### Supplementary Figure 9

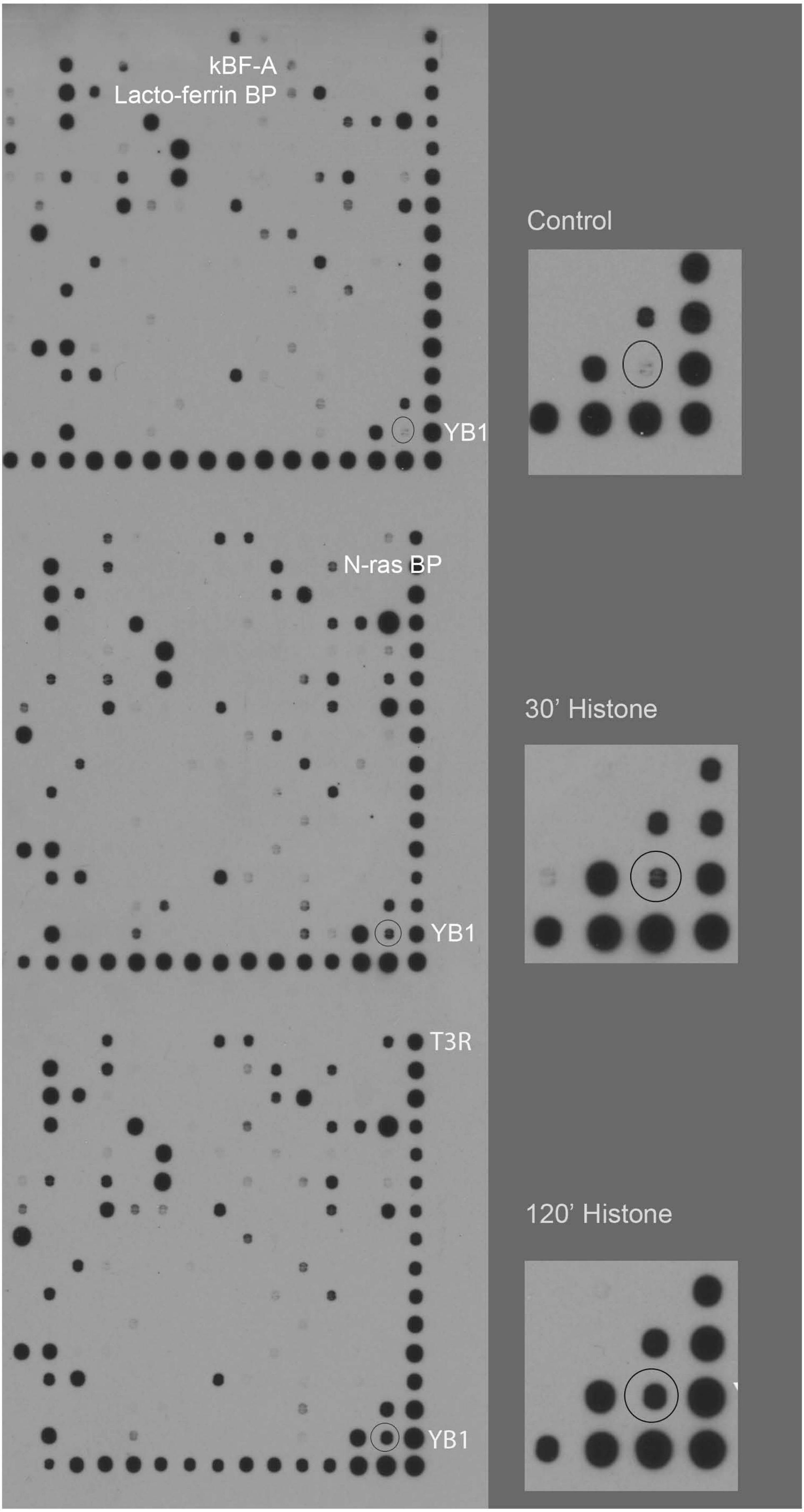

Supplementary Figure 10

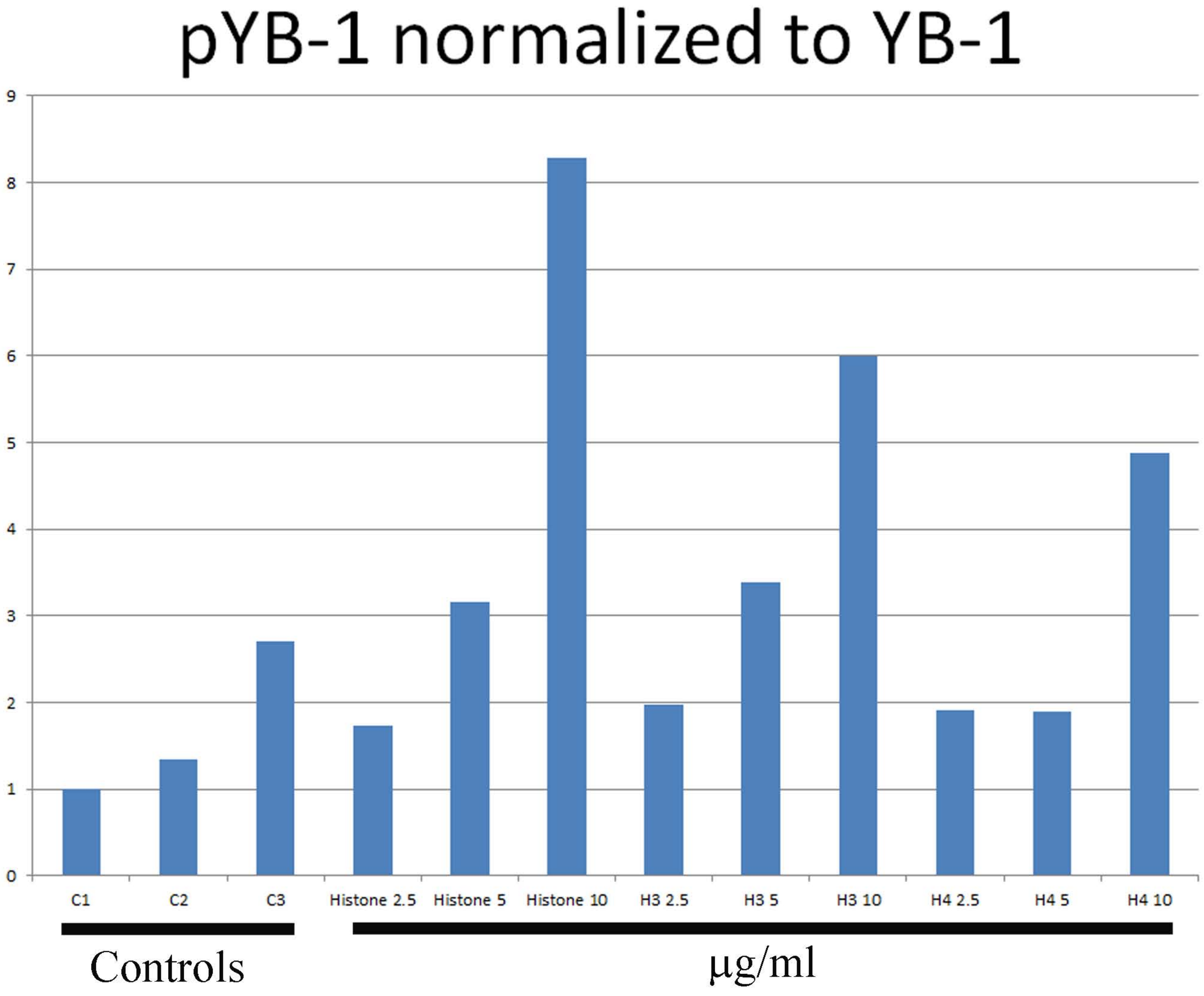

### Supplementary Figure 11

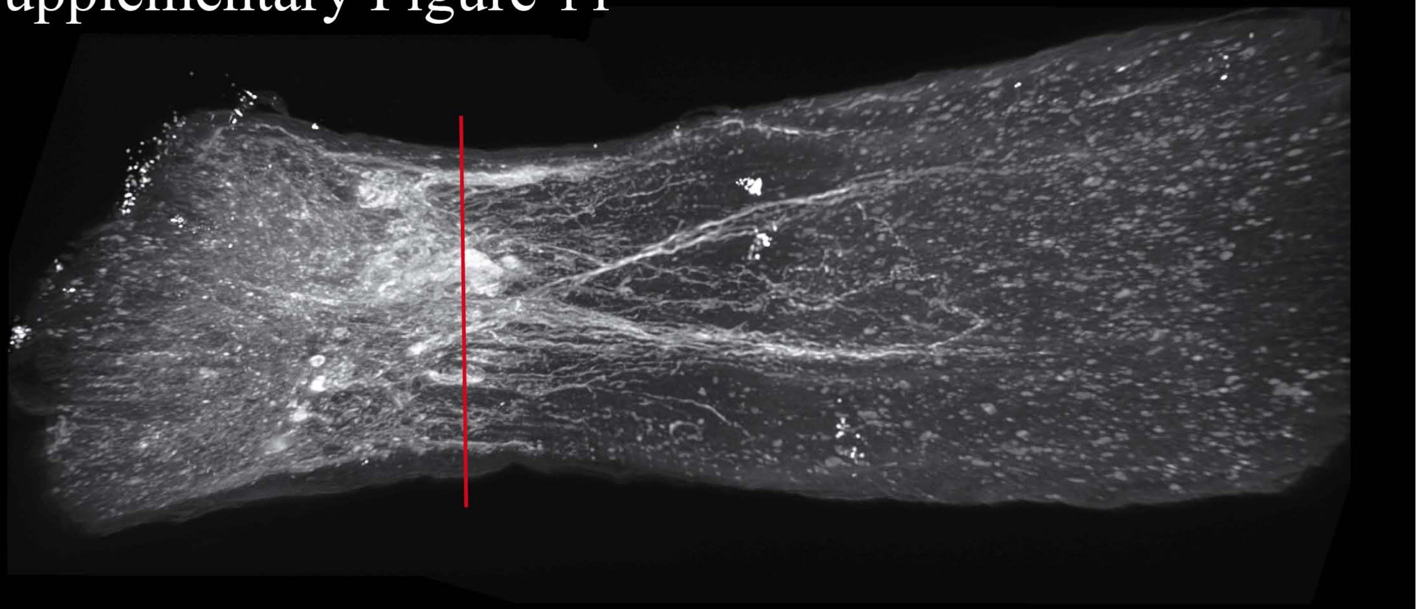

A.

B.

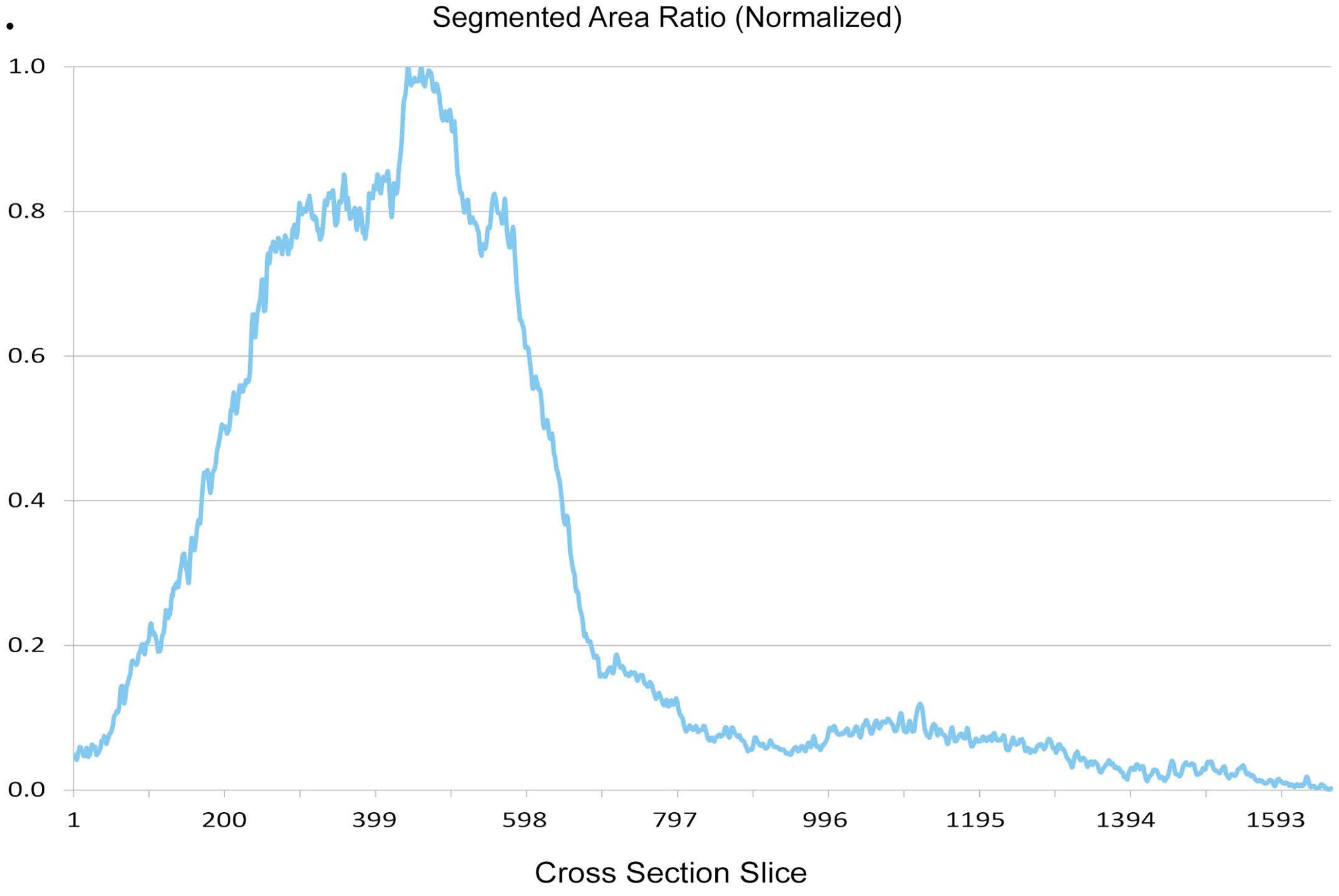

### Fig. S12

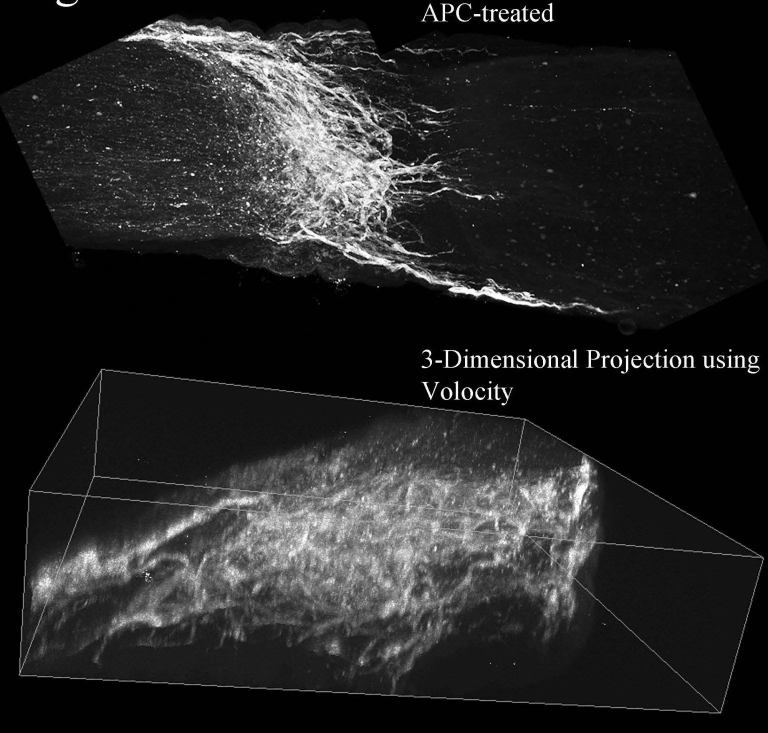

### Supplementary Figure 13

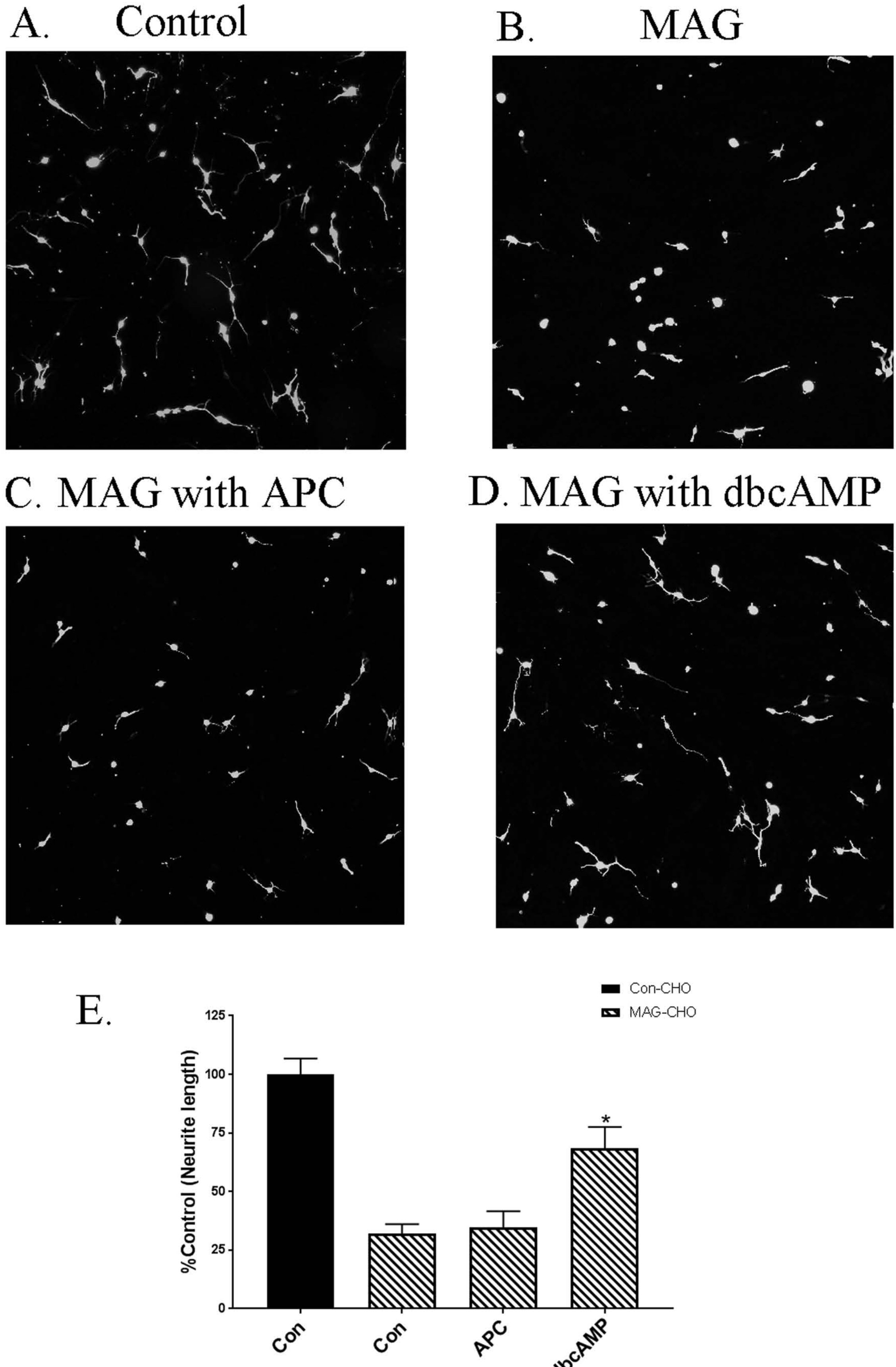
